# Allostery in Protein Tyrosine Phosphatases is Enabled by Divergent Dynamics

**DOI:** 10.1101/2023.07.23.550226

**Authors:** Colin L. Welsh, Lalima K. Madan

## Abstract

Dynamics-driven allostery provides important insights into the working mechanics of proteins, especially enzymes. In this study we employ this paradigm to answer a basic question: in enzyme superfamilies where the catalytic mechanism, active sites and protein fold are conserved, what accounts for the difference in the catalytic prowess of the individual members? We show that when subtle changes in sequence do not translate to changes in structure, they do translate to changes in dynamics. We use sequentially diverse PTP1B, TbPTP1, and YopH as the representatives of the conserved Protein Tyrosine Phosphatase (PTP) superfamily. Using amino acid network analysis of group behavior (community analysis) and influential node dominance on networks (eigenvector centrality), we explain the dynamic basis of catalytic variations seen between the three proteins. Importantly, we explain how a dynamics-based blueprint makes PTP1B amenable to allosteric control and how the same is abstracted in TbPTP1 and YopH.

## Introduction

Correlations between enzyme function and protein dynamics are gaining increasing interest as researchers explain biological function^1^, explore therapeutic opportunities^2^, and expand enzymes to new chemistries^3^. The classical view asserts the role of dynamics in influencing the stability of enzyme active sites^4^, participation in substrate binding^5^, and/or modulation of thermal stability of enzyme:substrate transition states^6^. A more recent view emphasizes the role of concerted amino acid networks in playing a role in catalysis and is inclusive to detailing distal sites that may have the ability to modulate enzyme function^5,7-10^. While the exact nature of protein dynamics’ contribution to enzyme catalysis can be debated^11,12^, it is apparent that cell signaling makes great use of protein dynamics, both enzymatically and otherwise^13^. Signaling enzymes, including protein kinases and protein phosphatases, are allosterically modulated, often exploiting dynamics-based components of their catalytic properties^8,14-18^. In this study we focus on identifying the dynamics-based distinctions between three Protein Tyrosine Phosphatases (PTPs) that share a near-identical active site but are varied in their catalytic rates.

PTPs are a superfamily of protein phosphatases^19,20^, that are principally responsible for the cleavage of phosphorylated tyrosine residues into tyrosine and inorganic phosphate. Of these, the Class I Classical PTPs (hereinafter referred to as just PTPs) are well-appreciated for their crucial roles in the modulation of human cell signaling^21,22^ and are targets of much scrutiny into their atomic details of function and regulation^23-25^. Recent work regarding the mobile loops that make up the canonical active site as well as the dynamic allostery inherent in Class I Classical PTPs highlights their prominence as models for investigating the impact of conformational dynamics on catalysis^15,16,24,26-35^. The conserved PTP domain is defined by a set of 10 conserved motifs^20^ (M1-M10) (Figure 1a); including 6 structural motifs (M2-M7) that form the core of the domain and 4 active site motifs (M1, M8-M10) that play a direct role in catalysis (Figure 1c). M1, the phosphotyrosine (pY)-binding loop or pY-loop, is defined by the sequence NxxKNRY/F and contains an aromatic residue (Tyr46 in PTP1B, Tyr51 in TbPTP1 and Phe229 in YopH) for binding the incoming pY substrate^36^. M8, the WPD-loop, is defined by the sequence (Y/F)xxWPDxGxP and houses the general acid catalyst aspartic acid (Asp181 in PTP1B, Asp199 in TbPTP1 and Asp356 in YopH) required for catalysis^37,38^. This loop can adopt multiple conformations such that open conformation(s) keep the enzyme in an inactive state, while the closed conformation is catalytically competent^39^. Rate of opening-closing of this loop is linked to PTP activity^29,31^ and blocking its closure by allosteric modulators is an effective strategy to inhibit PTP function^15,16,40^. M9, the Phosphatase-loop or P-loop, is defined by a 13 amino-acid sequence containing the HCx_5_R motif that is a signature of a PTP active site^41^. It contains the nucleophilic cysteine residue (Cys215 in PTP1B, Cys229 in TbPTP1 and Cys403 in YopH) that is stabilized in a deprotonated form for efficient catalysis^42,43^. Finally, M10, the Q-loop, is defined by the sequence QTxxQYxF that contains a glutamine residue (Gln262 in PTP1B, Gln275 in TbPTP1 and Gln446 in YopH) important for coordinating catalytic waters at the PTP active site^43,44^. The outer periphery of the PTP active site is marked by a conserved Glutamate^20^ (Glu115 in PTP1B, Glu126 in TbPTP1 and Glu290 in YopH) that resides in a sequence variable E-loop (Figure 2a). While the E-loop is not a part of the ten motifs detailed above, its functionality has garnered recent interest in PTP catalysis^15^ and redox regulation^45^.

**Figure 1.**
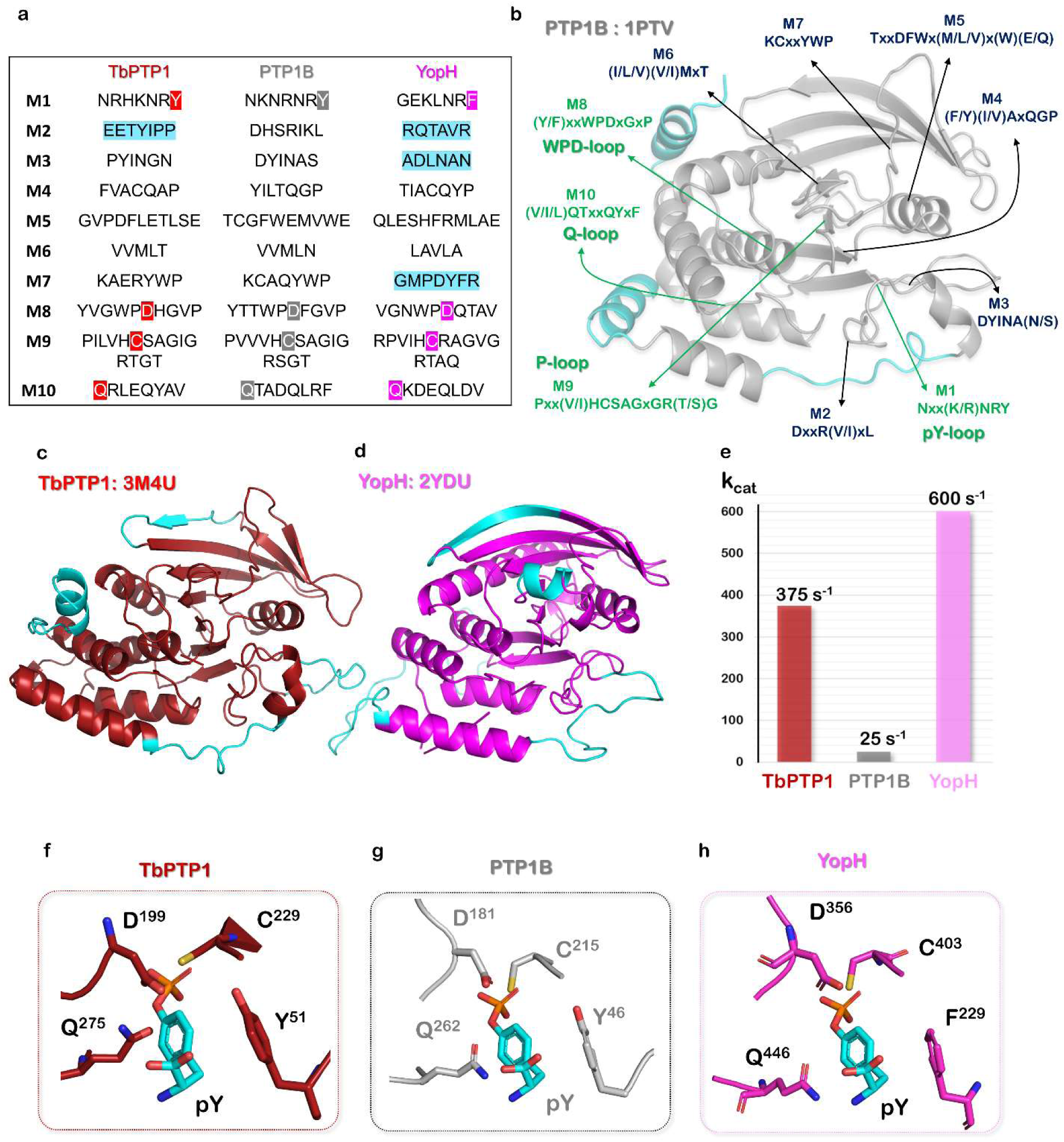
PTP domain organization and an overview of studied systems. (a) Sequences of the conserved PTP motifs as seen in TbPTP1, PTP1B and YopH. Catalytic residues are highlighted in red (TbPTP1), gray (PTP1B) and magenta (YopH). Sequence highlighted in Cyan show divergence from the canonical motif. (b) Ten conserved motifs mapped to the catalytic domain of PTP1B. PTP1B-specific structural regions are colored cyan. (c) Cartoon representation of the PTP domain of TbPTP1. Protein-specific regions as they diverge from the canonical PTP domain are colored in Cyan. (d) Cartoon representation of the PTP domain of YopH. Protein-specific regions as they diverge from the canonical PTP domain are colored in Cyan. (e) Kinetic properties of TbPTP1, PTP1B and YopH for hydrolyzing *para*-*NitroPhenyl Phosphate* (pNPP) (obtained from Ref ^56,62^). (F,G,H) Active site of the TbPTP1, PTP1B, and YopH binding a Phosphotyrosine residue, as seen in the starting structures of our pY-bound simulations.

**Figure 2.**
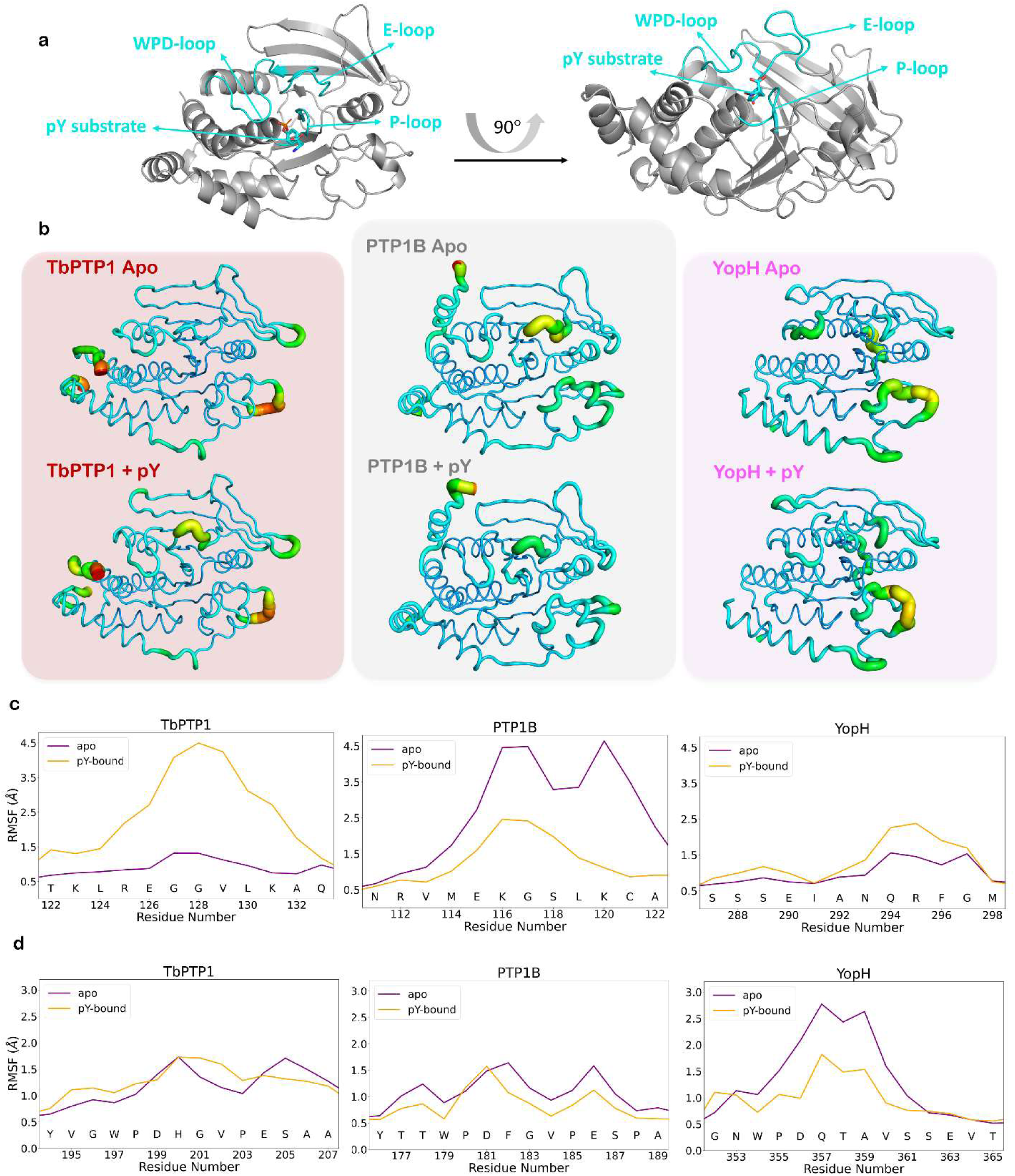
Essential dynamics and C*α* backbone fluctuations. (a) Structural arrangement of the active site loops. Loops in question are colored in cyan. (b) Putty representation of C*α* root-mean-squared fluctuations (RMSF). Increased ribbon diameter and red-shift through the VIBGYOR color spectrum indicates higher RMSF of that region. (c) Per-residue RMSF of residues in the E-loop for TbPTP1 (left), PTP1B (middle), and YopH (right). Apo state is shown in purple and liganded, pY-bound state is shown in orange for each enzyme. (d) Per-residue RMSF of residues in the WPD-loop with identical coloring and layout as in (c).

All PTPs use a conserved catalytic mechanism that proceeds in a two-step fashion (Figure S3A)^22,46,47^. In the first step, the tyrosine residue of the pY-loop interacts with the substrate pY. The sulfur atom of the deprotonated active site cysteine residue acts as a nucleophile and attacks the substrate phosphorous atom while the protonated aspartic acid residue of the WPD-loop donates it proton to the substrate tyrosine oxygen atom to create a better leaving group. This creates a cysteinyl-phosphate intermediate in the PTP active site, which is hydrolyzed in the second step. Here, the glutamine residue of the Q-loop coordinates catalytic water molecules while the now-deprotonated aspartate of the WPD-loop abstracts a proton, activating the water molecule. This nucleophile then attacks the phosphorous of the cysteinyl-phosphate intermediate, returning the enzyme to its initial state^48^. Given the conserved structure and mechanism of PTPs, it is easy to overlay details of molecular mechanics from one more popularly studied enzyme member to other less studied family members; diversity between members is often overlooked and even discounted while analyzing a family of enzymes. However, nature has created individual enzyme members with distinct catalytic prowess to suit the needs of the organism. How are these differences in catalytic activity achieved in a conserved active site that employs a conserved mechanism? Here, protein dynamics can provide the vital information to connect the gap between “sequence”, “structure”, and “function” paradigm.

In the presented study, we investigate the protein dynamics of the prototypical PTP1B, alongside its two most divergent PTP homolgues with a solved structure, YopH from *Yersinia pestis^49^* and TbPTP1 from *Trypanasoma brucei^50^* (Figure S2A). PTP1B is a mammalian enzyme, ubiquitously expressed in multiple tissues including the liver, skeletal muscle, adipose tissue, and the brain^51^. It is an important participant of the cellular signaling machinery and is most popularly known as a negative regulator of insulin and leptin signaling^51,52^. YopH is a secreted, highly-active PTP, that is a virulence determining factor for the parasitic *Yersinia pestis*^53^. YopH is an intriguing antibacterial drug target for curing plague and gastrointestinal syndromes^54,55^. Finally, TbPTP1 controls life cycle events and inhibits differentiation of African trypanosomes in mammalian hosts, preventing premature loss of immune evasion abilities^56,57^. PTP1B, TbPTP1, and YopH share a conserved PTP domain that superimposes with a cumulative RMSD of ∼2.5Å over their Cα backbone structures (Figures 1a, S1). TbPTP1 and YopH, being most divergent from PTP1B, have distinctions in sequence from PTP1B throughout their length. These include variations of the E-loop, between M6 and M7, as well as some positions on the WPD-loop (Figures 1a, S1B). These variations in YopH and TbPTP1 mirror the non-conserved nature of these same positions in the PTP family as garnered by a sequence alignment of the PFAM PTP family (Figure S2B,C). Some protein specific regions of the three include the secondary pY binding site in PTP1B^58^, the four trypanosome-specific motifs^59^, and the extended E-loop of YopH^60^ (Figure 1b). Despite the small differences, the active site of the three proteins is near-identical (Figure 1c) and uses the exact same two-step mechanism for hydrolysis of phosphotyrosines^26,57,61^.

The sharing of a highly conserved fold and catalytic motifs between the three proteins is somewhat incongruous with their vastly different catalytic activities (Figures 1b, S3B). YopH is the most active known PTPs with a *k_cat_* of 600s^-1^ towards *para-*nitrophenol phosphate (pNPP)^62^, a pan-PTP substrate that mimics a pY residue. TbPTP1 is also highly active with a *k_cat_* of 375s^-1^towards pNPP^59^; PTP1B, while quite active compared to other human PTPs, has a relatively low *k_cat_* of only 25s^-1^ for pNPP^62^. All three enzymes share a comparable affinity (*K_m_*) for pNPP (Figure S3B) as pNPP accesses their active sites without a contribution from protein surface residues. With a conserved fold, catalytic residues, and catalytic mechanism, this disparity between the activities of the three enzymes can only be explained by a difference in their dynamics. Our investigation into the evolutionarily diverse members of the PTP family, PTP1B (human), YopH (*Y. pestis*), and TbPTP1 (*T. brucei*) seeks to understand the fast-timescale motions of these enzymes using molecular dynamics (MD) simulations of both a pY-bound and ligand-free state (total six states). We analyze the essential dynamics of the six states using Principle Component Analysis (PCA)^63^ and their dynamics-based allostery by network analysis^64-66^. We characterize protein communities using the Girvan Newman approach^5,8,67^ for each state and identify the most influential nodes using Eigenvector centralities^68,69^. In comparing these three divergent enzymes, the similarities in dynamic features constitute the base requirements for PTP functionality, while the differences give rise to the variation in catalytic rates.

## Results and Discussion

### Essential Dynamics as seen in TbPTP1, PTP1B and YopH

The PTP active site is a phosphate binding P-loop nested between four loops (Figure 2a). Analysis of overall fluctuations of theses loops across the six systems reveal changes in backbone dynamics upon ligand binding (Figure 2b). Particularly, for PTP1B, the E-loop is seen to be highly mobile when the active site is unoccupied and does not house a ligand. These fluctuations decrease in the presence of pY but retain an RMSF of ∼2.5Å in the residues Lys116 and Gly117 (Figure 2c). However, TbPTP1 shows a opposite patten where the E-loop is rigid in the absence of the ligand but becomes mobile once pY binds its active site. YopH contains a rigid E-loop that adopts a α-helical conformation and shows minimum perturbations. The WPD-loop of all systems maintains a low level of mobility, with the exception of YopH in the unbound state (Figure 2b). In this system, the C_α_ fluctuations peak in the residues Gln357, Thr358, and Ala359 (Figure 2d). This increased mobility is likely due to the lack of the Pro residues in the sequence of the WPD-loop as seen in PTP1B and TbPTP1 (Figure 2d). Also, all systems show very high fluctuations in a region adjacent to the pY-loop (Figure 2b). However, due to their prevalence both with and without a ligand, it is likely that these fluctuations do not play a role in regulating catalysis. Instead, due to their proximity to the pY-loop, it is possible that these highly mobile regions are important for interactions with broader protein substrates in a region distinct from the substrate pY residue.

### Principal Component Analysis of Backbone Motions

In order to extract specific motions from the overall trajectory of each system, we performed PCA to analyze the motions seen in the RMSF (Figure 3, 4, S11). We analyzed the first five eigenvectors in-depth for each system, based on their contributions to the total variance and the RMSF explained by those motions (Figures S12-S16). Mapping the individual PCA modes back to the protein structures, we see the major motions of the protein backbone (Figure 3, 4, S11). In TbPTP1, there is a great deal of motion in the loop structurally adjacent to the pY-loop, but there seems to be little coherence to these motions, as most modes seem to be various motions of the loop. In the pY-bound state, we do see the motion of the E-loop previously noted (Figure 2b,c). This motion seems to occur mainly along one axis, in which the loop is either moving towards the active site or away from it, not side-to-side.

**Figure 3.**
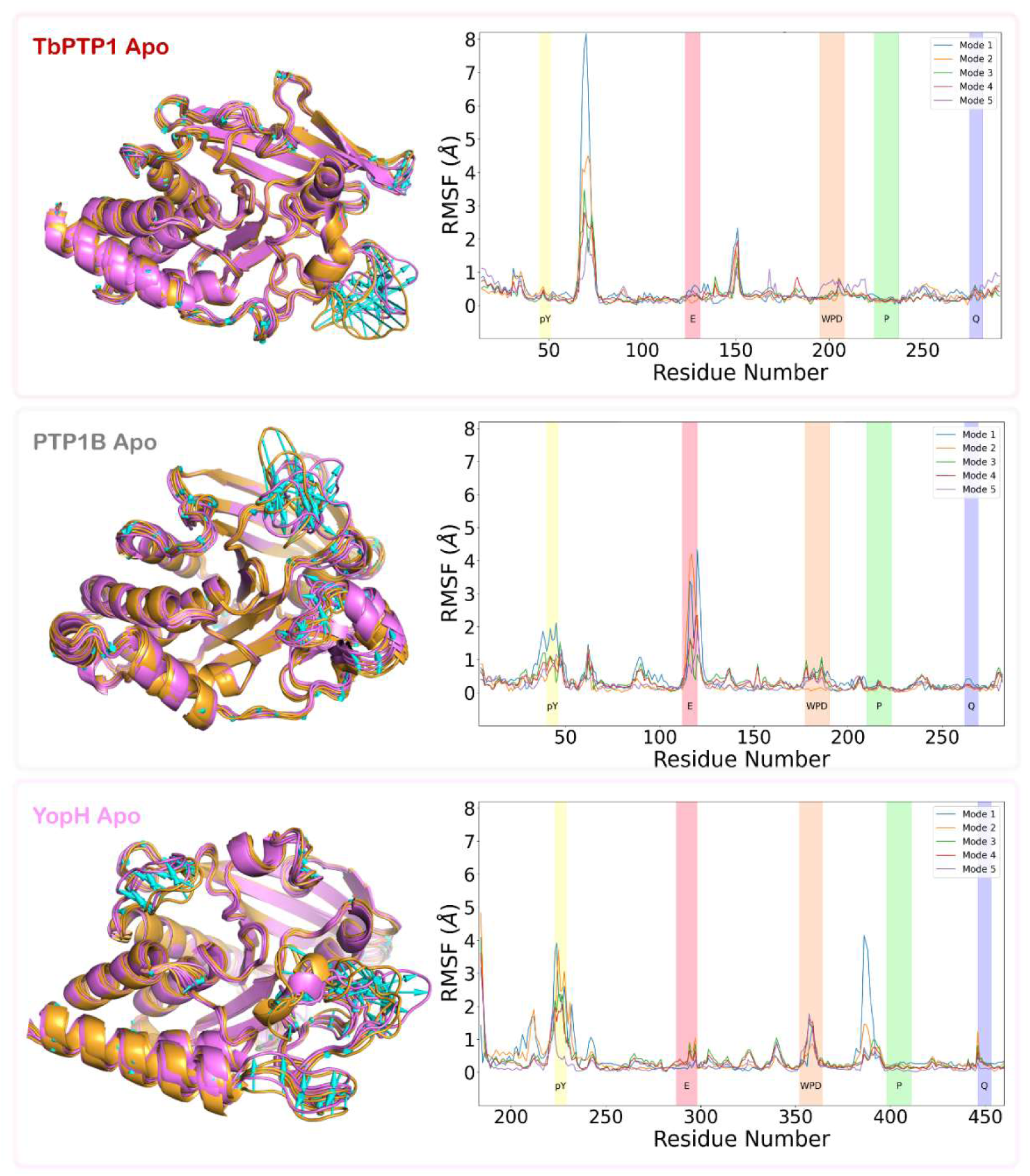
Principle Component Analysis of C*α* motions. Overlay of 5 first individual modes of motion as seen in the apo proteins (left column). Motions are shown for all three enzymes, TbPTP1 (top), PTP1B (middle), and YopH (bottom). Initial frames for each individual mode are in orange, the final frame in pink. Arrows in cyan show motions of C*α* atoms > 1.5Å. RMSF plots for the first 5 eigenvectors of PCA for TbPTP1 (top), PTP1B (middle), and YopH (bottom) systems are shown to the right. Catalytic motifs are highlighted: pY-loop (yellow), E-loop (red), WPD-loop (orange), P-loop (green), and Q-loop (blue).

**Figure 4.**
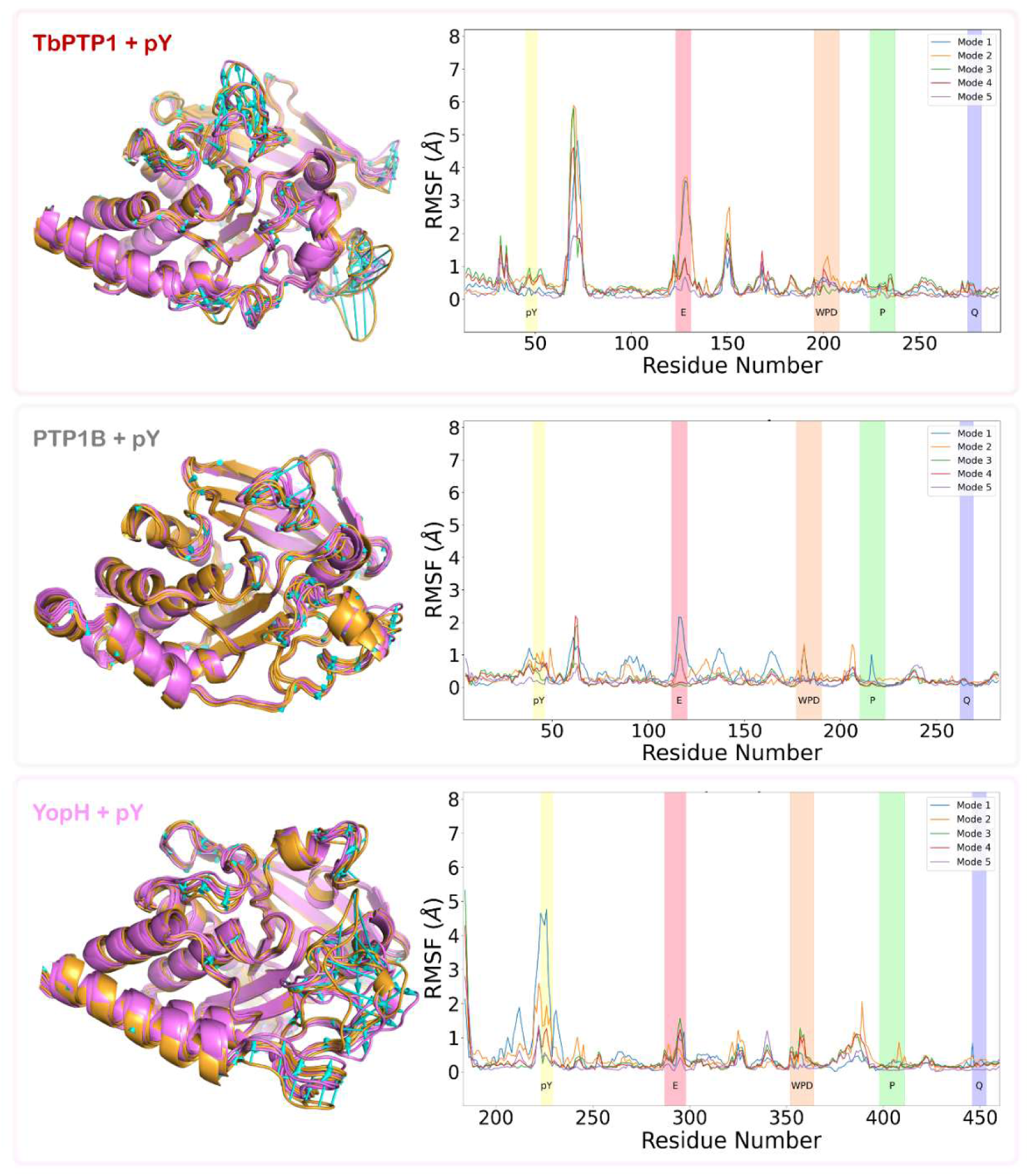
Principle Component Analysis of C*α* motions. Overlay of 5 first individual modes of motion as seen in the pY-bound protein complexes (left column). Motions are shown for all three enzymes, TbPTP1 (top), PTP1B (middle), and YopH (bottom). Initial frames for each individual mode are in orange, the final frame in pink. Arrows in cyan show motions of C*α* atoms > 1.5Å. RMSF plots for the first 5 eigenvectors of PCA for each system are shown to the right. Catalytic motifs are highlighted: pY-loop (yellow), E-loop (red), WPD-loop (orange), P-loop (green), and Q-loop (blue).

Comparatively, in the PTP1B apo state, the motions of the E-loop seem to be more random, moving along more than one axis. These motions are much less dramatic in the PTP1B pY-bound state. In fact, the motions represented by the various PCA modes are all very small in terms of displacement. This is likely due to the lower RMSD of the whole system (Figure S6) or the lower contributions of each individual PCA mode to the total variance (Figure S11), or a combination thereof.

In YopH, motions are concentrated in the WPD- and pY-loops (Figures 3, 4, S12-S16). There seems to be major deformations of the pY-loop in the unbound state that is reduced in the pY-bound state. This increased flexibility could partially explain the substrate promiscuity typical in YopH^70^. The WPD-loop motions are quite different between the two states, implying that these are not a general flexibility of the WPD-loop but instead offer some functionality to the enzyme.

To compare the two states of each enzyme, we calculated the cosine similarity index for each pair of the first five eigenvectors (Figure S16). This yields a value of 1 if perfectly parallel, -1 if perfectly antiparallel, and 0 if orthogonal. Due to the high dimensionality of these vectors, the highest value seen was 0.46 in TbPTP1. The increased similarity of the modes seen in TbPTP1 is likely due to the prevalence of major motions in the loop adjacent to the pY-loop in both states (Figures 3, 4, S12-S16). This implies that these motions are not strictly relevant to the binding or catalysis of the substrate pY itself; instead, due to the proximity of this loop and the pY-loop, it may be important for the binding of the broader protein substrate. Comparatively, in PTP1B and YopH, there is not a similar degree of similarity between the modes. The closest in between vector 5 in both PTP1B+pY and PTP1B-apo (Figures 3, 4, S12-S16). These modes capture inverse motions of the E-loop, with one moving away from the active site and the other moving toward the active site.

### Active Site Dynamics Rationalize Kinetic Properties

To assess differences in active site dynamics between the different enzymes, we analyzed the depth of the active site cleft, measuring the distance between the C_α_ atoms of the catalytic residues of the pY-loop (Tyr51 in TbPTP1, Tyr46 in PTP1B, and Phe229 in YopH), P-loop (Cys229 in TbPTP1, Cys215 in PTP1B and Cys403 in YopH) and the Q-loop (Gln275 in TbPTP1, Gln262 in PTP1B and Gln446 in YopH) over the length of MD simulation runs (Figures 5a, S17). In PTP1B, there is a single high-probability density conformation regardless of state. This is not the case in either TbPTP1 or YopH. In TbPTP1, a small conformational change is seen in the Q-C distance, showing a small deformation in the Q-loop. This is also seen in YopH, with the addition of the pY-loop motions increasing the Y-C distance in the unbound form.

**Figure 5.**
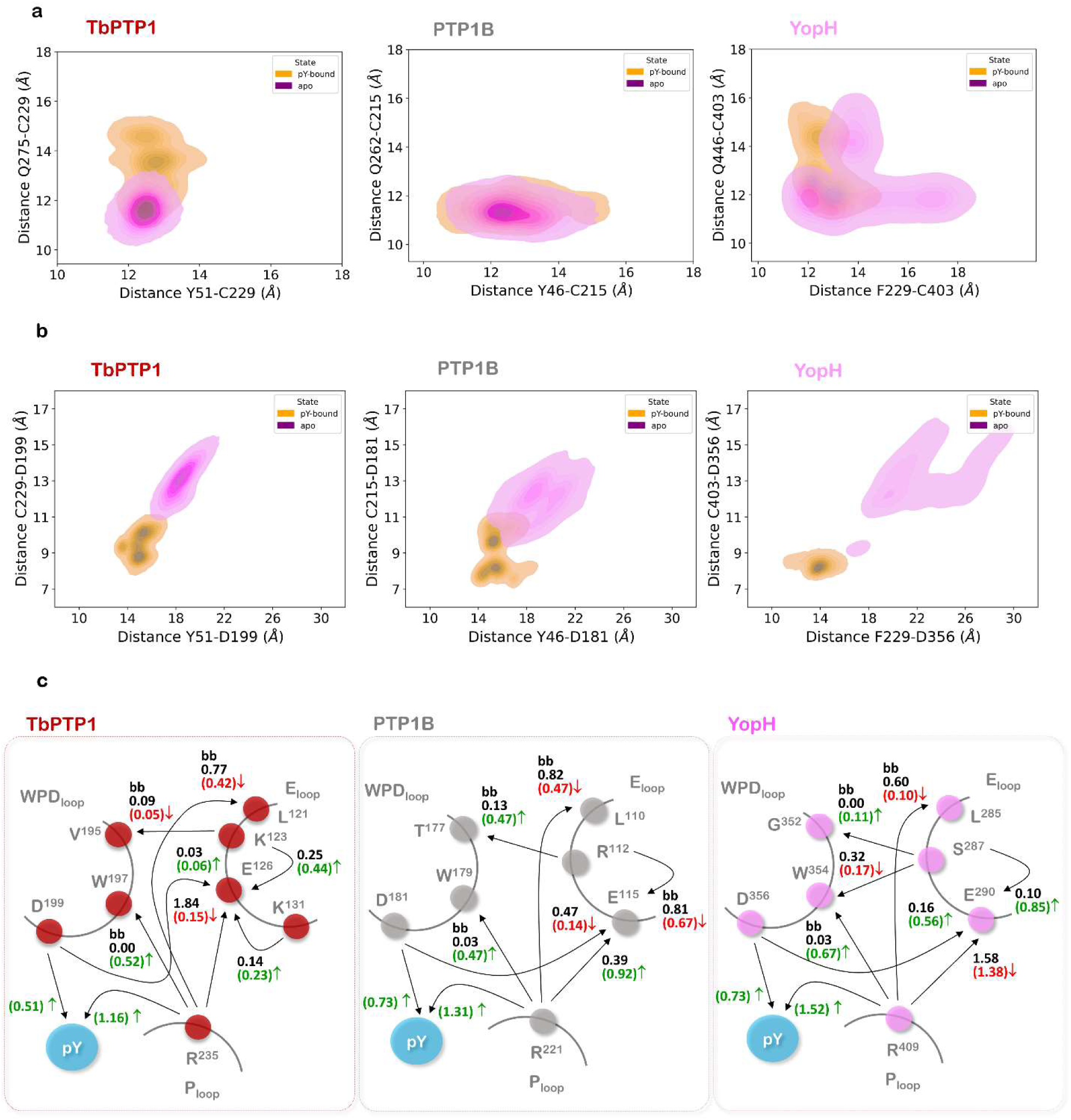
Active site dynamics. (a) Probability density between the C*α* atoms of residues on the Q-loop and P-loop (y-axis) and residues on the pY-loop and P-loop (x-axis) for TbPTP1 (left), PTP1B (middle), and YopH (right) in the unbound (purple) and pY-bound (orange) states. Darker colors indicate a higher density, or more frequent occurrence, of that pair of distances. (b) Identical to (a), except measuring the distance between the C*α* atoms of residues of the P-loop and WPD-loop (y-axis) and of the pY-loop and WPD-loop (x-axis). (c) Hydrogen bond interactions between residues of the P-, WPD-, and E-loops for TbPTP1 (left, red), PTP1B (middle, gray), and YopH (right, pink). The arrow points from hydrogen bond donor to hydrogen bond acceptor, with “bb” indicating the involvement of backbone atoms. The top, black number indicates the proportion of frames in which that hydrogen bond is present in the unbound form. The bottom, colored number indicates the proportion of frames in which that hydrogen is present in the bound form, indicating either an increase (green) or decrease (red) in proportion from the unbound state. Note that many residues can have more than one donor or acceptor atom, thus the proportion of frames can be greater than 1.

The WPD-loop dynamics show a much more distinct conformational change between states (Figure 5b). Comparing unbound states, TbPTP1 adopts a single high-density WPD-loop open conformation, unlike PTP1B, which samples two similar conformations, or YopH, which shows diffuse density, implying a mobile WPD-loop in the apo form. The additional density seen in the top-right corner of the YopH-apo plot is due to the deformation of the pY-loop, not the hyper-open conformation mentioned in literature^15^. Interestingly, the WPD-loop of YopH adopts a single high-density conformation in the pY-bound state, whereas the other two enzymes adopt two high-density conformations, as well as other low-density conformations.

When considering these conformations in the context of the two distinct reaction steps (Figure S3A), they provide insight into the difference in catalytic rates between PTPs. The rates for the two steps, *k_cleavage_* and *k_hydrolysis_*, differ between the enzymes PTP1B and YopH with the substrate pNPP. In PTP1B, *k_cleavage_* is 270s^-1^ and *k_hydrolysis_* is 28s^-1^ from one study^27^. Comparatively, YopH was found to have *k_cleavage_* and *k_hydrolysis_* values of 343s^-1^ and 76s^-1^, respectively^71^. While no such mechanistic studies exist for TbPTP1, the *k_cat_* value for the overall reaction lies in-between PTP1B and YopH (Figure 1c). The conformations of TbPTP1’s catalytic motifs mirror characteristics of both PTP1B and YopH, in which the Q-loop behaves similarly in TbPTP1 and YopH (Figure 5a), but the WPD-loop of TbPTP1 behaves more akin to PTP1B (Figure 5b). The alternative conformation of the Q-loop in YopH and TbPTP1 could allow for increased interactions with the solvent and the recruitment of the catalytic water, increasing the rate of *k_hydrolysis_*, whereas the alternative conformations of the WPD-loop in PTP1B and TbPTP1 may indicate closed, but inactive, conformations that reduce the rate of *k_cleavage_*.

### Hydrogen-bond dynamics reveals the conserved role of Arg in the Cx_5_R motif

We analyzed the hydrogen bond dynamics at the interface of the WPD-, P-, and E-loops due to the highly polar nature of PTP active sites and to examine the proposed role of the E-loop in WPD-loop dynamics (Figure 5c). This analysis considers a hydrogen bond present between two atoms less than 3Å apart with a donor atom-hydrogen atom-acceptor atom angle greater than 135°. When analyzing the prevalence of hydrogen bonds in the active site, through determining the proportion of frames in the trajectory where the criteria is met, we identified dramatic changes in the hydrogen bond network that occur after ligand binding. Due to the possibility of having more than one donor-acceptor pair per residue (i.e., the guanidinium group on arginine and the phosphate group on phosphotyrosine), it is possible that the proportion of frames where an H-bond is present is greater than one.

This analysis highlighted the importance of the P-loop Arginine residue (Arg235 in TbPTP1, Arg 221 in PTP1B and Arg405 in YopH) that is a part of the conserved HCx_5_R active site motif. In all three enzymes in the unbound state, this Arg makes a prevalent H-bond with the backbone of a Leu at the base of the E-loop (Leu121 in TbPTP1, Leu110 in PTP1B and Leu285 in YopH) (Figure 5c). This Arg also makes H-bonds with the conserved Glu of the E-loop (Glu126 in TbPTP1, Glu115 in PTP1B and Glu290 in YopH), but the prevalence of this interaction is much higher in YopH (1.58) and TbPTP1 (1.84) when compared to PTP1B (0.39) (Figure 5c). This interaction seems important for the mobility of the E-loop. Upon ligand binding, the proportions for this interaction change to 0.92, 1.38, and 0.15 for PTP1B, YopH, and TbPTP1, respectively, implying an inverse relationship between the prevalence of this interaction and the mobility of the E-loop. When considering this alongside the H-bonds made between this Arg and the substrate pY, it is possible that the E-loop interaction is important for R221 to adopt a conformation suitable for ligand binding. Indeed, the distribution of its sidechain χ-angles shift in the presence of the ligand, particularly in χ_3_ and χ_4_ (Figure S18). A key change that occurs in the presence of the ligand pY is the H-bond between the conserved Arg and the backbone Trp of the WPD-loop (Trp197 in TbPTP1, Trp179 in PTP1B and Trp354 in YopH) (Figure 5c). This increase in prevalence is seen in all three enzymes in the pY-bound state, and when compared with the H-bonds present between (PTP1B Numbering) Arg221 and Leu110 in the unbound state, suggests a “switch” mechanism in which a ligand-induced conformational change of Arg221 creates an H-bond with Trp179, aiding in the closure of the WPD-loop in all three enzymes. Further interactions are seen between many residues on the E-, WPD-, and P-loops, but differ in trend and prevalence between the enzymes. These residues surely play a role in determining the rate of closure or flexibility of the WPD-loop for that enzyme, but it is likely that these enzymes use a conserved mechanism for the basal WPD-loop closure.

### PTP Active Sites of TbPTP1/PTP1B/YopH partition into distinct “Communities”

We employed a network-based analysis to gain a more detailed view of the dynamics of the enzymes^8^. From a cross-correlation-based network that separately determines the correlation for a residue’s main chain and side chain, we utilized the Girvan-Newman algorithm^72^ to detect communities of residues. In a given community, the residues (or portions of residues) within that community will be highly interconnected with more correlated motion compared to residues outside of that community. Detected communities are colored according to the catalytic residues contained within that community or its general structural region (Figure 6). The community containing prominent residues of the WPD-loop are colored orange, the Q-loop purple, E-loop dark red, pY-loop violet, and P-loop yellow. The number of communities detected are somewhat consistent among the six systems, with TbPTP1 containing 11 communities in the unbound and 16 communities in the bound form, PTP1B-apo and pY-bound containing 15 and 14 communities respectively, and YopH-apo contains 14 communities whereas the bound form contains 11 communities (Figure 6).

**Figure 6.**
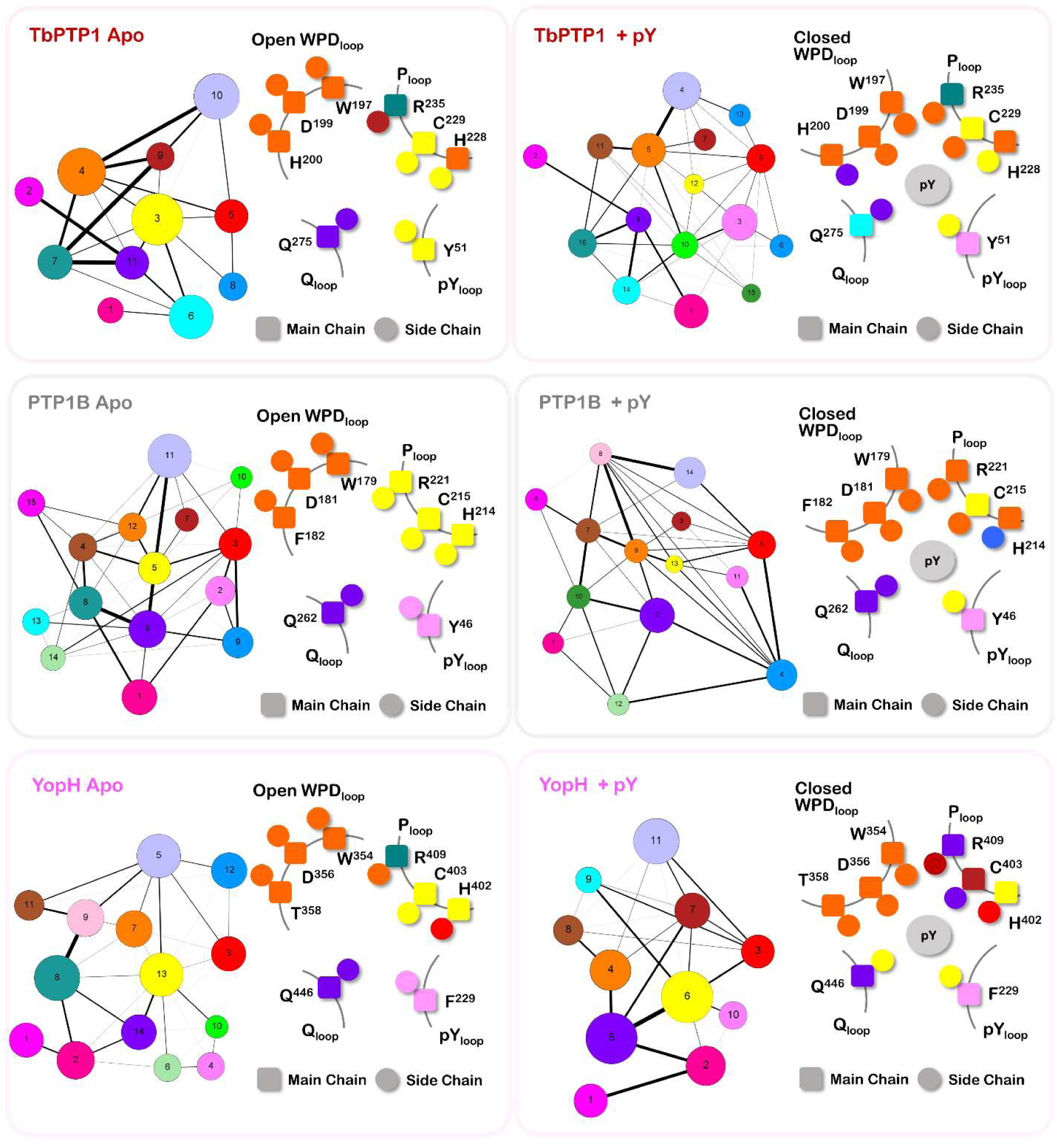
Girvan-Newman community analysis of TbPTP1(top), PTP1B (middle) and YopH (bottom) in their apo (left) and pY-bound (right) states. In each panel, the overall community distribution is shown (left of each), where the size of each community corresponds to the number of nodes it contains. Edges connected the communities are weighted (thickness of the connection) based on betweenness. The distribution of active site nodes is also shown (right of each), with the main chain being represented by squares and the side chains represented by circles.

When investigating the active site in more specific detail, several general trends emerge. Prominent residues in the WPD-loop remain in their own community in almost every enzyme and state, (apart from TbPTP1+pY, where the sidechain of His200 lies in the same community as the Q-loop (Figure 6). The Q-loop also seems to be mostly self-contained, except in TbPTP1+pY system, where the main chain of Gln275 resides in the α5 helix community (Figures 6, S19), and the YopH+pY system, where the sidechain of Gln446 resides in the P-loop community (Figures 6, S21).

The partitioning of notable P-loop residues to several different communities in almost every system implies that the dynamics of the P-loop are not distinct from its surroundings. This is best seen in the closed, substrate-bound state of YopH (Figures 6, S21), in which the main and side chains of the three conserved residues of the HCx_5_R motif are almost all in different communities. The implication of this partitioning is that the conserved positions of the P-loop serve as a “communication hub”, reading the dynamics of the surrounding structural features of the active site. This phenomenon concerning the P-loop is seen in almost all systems, regardless of state, except for PTP1B-apo.

The unbound PTP1B system maintains each active site feature in its own community (Figure 6). This lack of intercorrelation in the unbound state explains the lower activity seen in PTP1B, as the enzyme must dramatically alter the dynamics within the active site to allow for catalysis. Comparatively, while the specific communities may not remain the same, there is much more inter-loop communication in both TbPTP1 and YopH in both the apo and pY-bound states.

### Influential Nodes within the Dynamic Network

From our dynamic residue network, we were able to identify important residues through the calculation of eigenvector centrality (EC). This measure of a node’s influence on the network revealed similar residues among our six systems, with several notable variations (Figure 7). Prominently, catalytic residues are almost entirely absent in the set of influential nodes, with only the main chain of the catalytic cysteine (Cys229 in TbPTP1, Cys215 in PTP1B and Cys403 in YopH) having an EC > 0.10 in the TbPTP1-apo (Figures 7, S19, S22), PTP1B+pY (Figures 7, S20, S25), and YopH-apo (Figure 7, S21, S26) systems. This observation supports a dichotomy for the purpose of conserved residues, in which certain positions are important either for catalysis or for communication.

**Figure 7.**
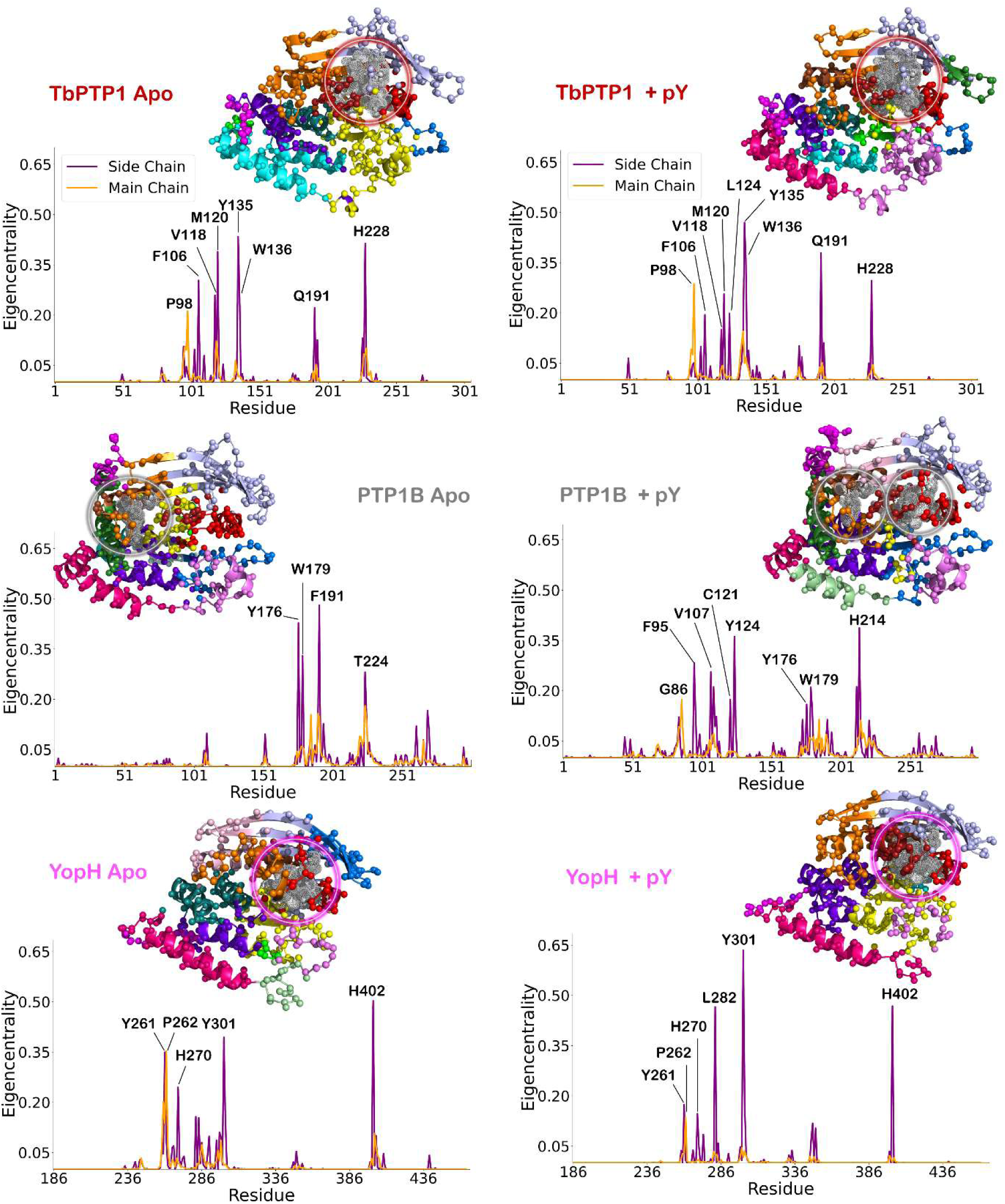
Influential nodes within the network for TbPTP1(top), PTP1B (middle) and YopH (bottom) in their apo (left) and pY-bound (right) states. The eigenvector centrality of side chain nodes are in purple and main chain nodes in orange. Notable residues as labeled on the plot are shown on the structure (showing Girvan Newman communities as in Figure 6) and are highlighted by circles. Also see Figures S19-S21.

We did not identify any residues in the variable region of the E-loop with EC > 0.10. However, in the more active TbPTP1 and YopH, residues in M6 (TbPTP1: 118-122; PTP1B: 107-111; YopH: 282-286) and M7 (TbPTP1: 131-137; PTP1B: 120-126; YopH: 297-303) that flank the E-loop showed much greater EC values. The side chains of residues Leu282 and Tyr310 in YopH showed high influence both with and without a ligand (Figures 7, S21, S26, S27); TbPTP1 in both states also shows high influence in the side chains of the corresponding Val118 and Tyr136, but also in the side chain of Met120 (Figures 7, S19, S22, S23). Notably, in the less active PTP1B, the unbound state showed M6 and M7 having very little influence on the dynamic network. This shifted in the presence of a ligand, in which the side chains of PTP1B’s corresponding residues Val107, Tyr124, and Met109 appear (Figures 7, S20, S24, S25).

Another interesting feature of PTP1B is the outsized influence of the WPD-loop in the unbound/apo state (Figures 7, S20, S24) when compared to all other systems, including the PTP1B+pY system. In the PTP1B-apo system, the only residues with main- or sidechain EC > 0.1 are Tyr176, Trp179, Pro185, Ser190, Phe191, Phe194, Thr224, Phe225, and Phe269. These residues all lie at the interface of the WPD-loop and the rest of the active site, specifically the α3, α4, and α6 helices. In stark contrast, no WPD-loop or adjacent residues show large influence in neither state of TbPTP1 nor YopH (Figures 7, S19-S27). The unique influence of PTP1B’s WPD-loop without a ligand certainly decreases in the presence of pY, but its influence does not disappear. In the closed form, there is a shift in which residues have an EC > 0.1, but many remain, specifically Tyr176, Trp179, Pro185, and Phe191, while the residues Pro180 and Arg221 appear (Figures 7, S20, S24).

In the network map of the PTP1B+pY system, the clusters of high-EC nodes seem to split into two separate groups (Figures S20, S25). One contains the residues of Motifs 6 and 7 that is consistent with YopH and TbPTP1, and a separate cluster made up of the now less influential WPD-loop residues, showing distinct dynamic mechanisms at work in PTP1B. This again contrasts the highly active YopH and TbPTP1, whose network maps show single clusters of high-EC nodes (Figures S23, S27).

## Discussion

As crucial regulators of cell signaling, PTPs have been the subject of extensive research to characterize their functionality and activity. However, this work has illuminated the seemingly paradoxical dichotomy of vastly different enzymatic activity with a highly conserved fold, which posits the dynamic motions of PTPs as the mechanism for divergent catalytic rates. As such, the dynamics of PTP1B and YopH have been the subject of numerous studies, both in the context of catalytic activity^15,26-32^ and in PTP1B’s allosteric regulation^16,24,33-35^, which likely exploits the fast-timescale motions of PTPs.

In this work, we examine the changes in protein dynamics that occur between the unbound and liganded, precatalytic states in three evolutionarily diverse PTPs. We explore the changes in backbone RMSF (Figure 2) seen in active site features, such as the WPD-, pY-, and E-loops, that occur upon substrate binding. We note an increased flexibility of the pY-loop of YopH, explaining the enhanced substrate promiscuity of that enzyme, as well as increased mobility of the WPD-loop in the open state. The mobility of the E-loop depending on state is contrasted between PTP1B and TbPTP1, where the human PTP1B is more mobile without a ligand, whereas the *T. brucei* TbPTP1 E-loop is rigid in the unbound state, mirroring the more active YopH.

Principal component analysis and eigenvector cosine similarity (Figures 3, 4, S11-S16) of these backbone motions reveal the distinct motions of the two states of each enzyme. The dramatic mobility of the TbPTP1 insert spatially adjacent to the pY-loop in both states, and the similarity of motion between states, implies its role is unrelated to catalysis. While both PTP1B-apo and TbPTP1+pY both saw mobility in their E-loops, these motions differ substantially in their directionality. Whereas the PTP1B E-loop moves in a multitude of directions, the TbPTP1 E-loop moves along a single axis.

Through analysis of the probability distributions of active site distances (Figure 5a,b), we detected an alternative conformation of the Q-loop in the more active PTPs, possibly creating the increased rate of the second catalytic step. Relevant to the rate of the first step, we saw multiple high-density conformations for the WPD-loop in PTP1B and TbPTP1, explaining the depressed catalytic rate compared to YopH. We also analyzed the hydrogen bonding network (Figure 5c) of the highly basic PTP active site, identifying a hydrogen bond switch in the Arg221-Leu110-Glu115/Arg221-Trp179-Glu115 interaction (PTP1B numbering) that selectively stabilizes either the WPD- or E-loops.

Community detection from the dynamic cross-correlation network (Figure 6) elucidated the differential distribution of communities in the PTP active site. The active site features of YopH and TbPTP1, the more active enzymes, do not segregate into entirely distinct communities and instead have several overlaps between communities and structural elements. This contrasts with PTP1B, where the community distribution of the active site is dependent on the sequence or structure of the active site loops in the unbound state. In the active form, this distribution changes to mirror the more active enzymes more closely, in that there is more overlap in communities between the loops.

The identification of influential nodes through the eigenvector centrality (Figure 7) illuminates potential regulatory mechanisms that have evolved in mammalian PTPs. In the parasitic PTPs TbPTP1 and YopH, the distribution of influential nodes remains mostly consistent between the unbound and bound states. Particularly, residues in M6 and M7, which flank the E-loop, and the histidine residue from the HCX_5_R motif show high eigenvector centrality. This contrasts PTP1B, which shows different patterns of influential nodes between the two states. In the unbound form, residues in and adjacent to the WPD-loop showed high influence. With the pY ligand, the pattern of influential nodes appears more similar to YopH and TbPTP1.

Taken altogether, the dynamic landscapes of these three enzymes mirror the evolutionary philosophy/ biological-need behind their function. PTP1B is a crucial signaling regulator in human cells, modulating the action of kinases such as Janus kinase (JAK) and the insulin receptor (IR), and as such, is heavily regulated through a myriad of mechanisms^73^. On the opposite end of the spectrum is YopH, a secreted virulence factor from the plague-causing *Yersinia pestis^70^*, whose high activity and broad substrate promiscuity benefits its expressing organism^71,74^. TbPTP1 lies somewhere in the middle of this spectrum. As an enzyme important for preventing premature life cycle progression of *Trypanasoma brucei*^50,59,75^, TbPTP1 must be highly active until it is no longer required, at which point mechanisms that are less nuanced than their mammalian counterparts can regulate this enzyme^76^. Our studies show that the structural dynamics of these enzymes mirror their responsibilities of being critical catalysts with regulatory control. Motions or interactions that are precisely conserved across the three PTPs likely enable basal catalysis; less similar processes create the differences in catalytic activity, and the divergent motions determine how these enzymes are regulated in their specific cellular environments. Future work should investigate dynamics with these concepts in mind, to determine precise mechanisms for loop closure and transition state stabilization between PTPs, but also to determine how human PTPs could be targeted therapeutically through exploiting inherent dynamic processes.

## Materials and Methods

### Multiple Sequence Alignment and Phylogenetic Analysis

The sequence conservation of PTPs was assessed through a multiple sequence alignment of the PFAM^77^ family entry y_phosphatase (ID: PF00102) with the exclusion of sequences <200 amino acids to exclude PRL phosphatases, which yielded ∼42,000 sequences. The database was aligned using MUSCLE^78^ with the Super5 algorithm. The alignment was analyzed using Jalview^79^, which also calculated the property conservation^80^ and percent consensus at each residue position. The sequences of solved structures of PTPs were also attained from the PFAM database and aligned with MUSCLE, upon which a phylogeny was built using the Phylogeny.fr server^81^ with the BioNJ distance algorithm^82^ and 100 bootstrap steps.

### System Preparation for Molecular Dynamics

This study investigated three enzymes: *Trypanasoma brucei* TbPTP1 (PDB ID: 3M4U)^50^, human PTP1B (PDB ID: 1PTV)^58^, and *Yersinia pestis* YopH (PDB ID: 2YDU)^49^. Missing loop regions from these structures were built using MODELLER^83^. For PTP1B, the active site C215S mutation present in the crystal structure was reverted to cysteine. For pY-bound systems, the pY ligand was aligned in the pocket using the pose present in the PTP1B crystal structure (PDB ID: 1PTV) prior to initial minimization, which was done in the gas phase for 2000 steps of steepest decent (SD) and 200 steps of conjugate gradient (CG) minimization in UCSF Chimera^84^. Model quality following minimization was assessed using PROCHECK^85^. Protonation states for most residues were assigned using PROPKA^86,87^, with the exception of the catalytic and active site residues aspartic acid (protonated) (TbPTP1: D199; PTP1B: D181; YopH: D356), cysteine (deprotonated) (TbPTP1: C229; PTP1B: C215; YopH: C403), and histidine (doubly-protonated) (TbPTP1: H228; PTP1B: H214; Yoph: H402) in order to simulate the enzymes in a catalytically competent state according to biochemical requirements and previous studies^45^.

### Molecular Dynamics Simulations

All MD simulations were performed using AMBER20^88^ with the ff14SB force field^89^. The phosphorylated tyrosine (pY) ligand was handled by the phosaa14SB force field ^90^. Hydrogen bonds were constrained using the SHAKE algorithm^91^ and the Particle mesh Ewald summation was used to handle long-range electrostatics^92^. Solvation was performed using the TIP3P^93^ water model with 12 Å padding from the periodic boundary. A constant temperature of 300 K was maintained using Langevin dynamics^94^ and constant pressure maintained by Berendson’s barostat^95^.

Minimization following solvation was performed first on the solvent and ions only by restraining the solute for 2700 steps of SD and 300 steps of CG. This restraint was gradually relaxed in phases of 50 kcal mol^-1^ Å^-2^, 20 kcal mol^-1^ Å^-2^, and 5 kcal mol^-1^ Å^-2^ for 8000 SD steps and 2000 CG steps. The system was then gradually heated from 0 K to 300 K in six steps of 50 K with a solute restraint of 5 kcal mol^-1^ Å^-2^ under the NVT ensemble. Further minimization was then performed by reducing the solute restraint to 2, 0.1 and 0.05 kcal mol^-1^ Å^-2^ for 8000 SD and 2000 CG steps per restraint value under the NVE ensemble. Following this, the system is equilibrated in NPT with restraints of 0.5, 0.1, 0.04, and 0.01 kcal mol^-1^ Å^-2^ for 50, 100, 400, and 400 ps. Finally, the system undergoes a 10ns unrestrained run with a 2fs timestep before the 250ns production run. 100ns was discarded from the beginning of each run to avoid equilibration artifacts based on analysis of structural factors (Figures S3-S8). A total of 4 replicates was run for all six systems (three enzymes, with and without pY ligand), yielding a total of 600ns data for each system.

### Trajectory Analysis

All analysis of MD trajectories was performed in CPPTRAJ^96^ unless stated otherwise. Standard structural factors of root-mean-squared deviation (RMSD), radius of gyration (R_G_), and solvent-accessible surface area (SASA) were analyzed to determine the equilibration of each system (Figures S4-S8). Principal component analysis (PCA) was performed on the C_α_ atoms of the core PTP domain of each system, defined by the residues 14-291 (TbPTP1), 5-282 (PTP1B), and 184-391 (YopH). Eigenvector similarity was determined in Python by calculating the cosine similarity between vectors *A* and *B*:

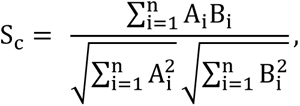

where *A_i_* and *B_i_* are components of vector *A* and *B*, respectively. Structure figures were made using PyMOL, graphs were made using the python based visualization packages MatPlotlib^97^ and seaborn^98^.

### Network construction and Community analysis

The construction of the dynamic network was performed using the networkView^99^ plugin of VMD^100^ as has been reported elsewhere^64-66^. The network is described as a set of nodes and edges, the nodes consisting of all amino acid residues split into two: the α node, containing the C_α_, C, O, and N atoms of the main chain, and the β node, containing all heavy atoms of the sidechain. An edge is created between every pair of nodes if any heavy atoms of the two nodes are within a contact distance 4.5 Å. The edge weight, *w_ij_*, is defined as a transformation of the dynamic cross correlation: *w_ij_* = − ln|*C_ij_*|. A final restriction that no edge is created between nodes belonging to the same residue is applied to the adjacency matrix before further analysis. This weighted adjacency matrix is used in a Python NetworkX^101^ implementation of the ForceAtlas2 algorithm^102^ for network visualization and layout.

Community analysis was performed to detect groups of nodes that are highly interconnected and intercorrelated among each other, and not with the nodes of other groups. The Girvan-Newman algorithm^72^ was employed to determine the community structures within the network. This algorithm functions in a top-down manner, iteratively removing edges that have the highest edge betweenness, which is the number of shortest paths between every pair of nodes that pass through a given edge. As edges are removed, the network is remembered whenever the number of communities, or groups of nodes that are disconnected from the rest of the network, increases. Once every edge is removed, the assignment of communities to nodes is selected that maximizes modularity, a measure of the probability difference of intra- and inter-community edges. Graphs were made using the open source package Gephi^103^.

Eigenvector centrality (EC) was calculated to identify influential nodes within the network. EC serves as a powerful method to integrate direct centrality (measure of direct connections) and information flow through the nodes^104^. This method is recently gaining traction for defining protein allosteric networks^68^, protein-ligand interaction^105^ and locating influential nodes that determine protein stability and function ^106,107^. EC for a node *x* is defined as:

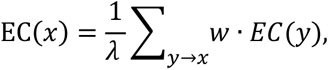

where *w* is the edge weight, *y* → *x* indicates all nodes *y* which are adjacent to *x* in a graph *G* = (*V*, *E*). This can be rewritten as *λe* = *Ae*, in which *A* is the weighted adjacency matrix, *λ* is the largest eigenvalue of *A*, and *e* is the EC of all nodes, such that the *i*th value of *e* is the EC of the *i*th node in *V*^108^. The EC of every node *x* is hence a measure of long-range interactions and connected to all other residues of the protein. Due to the recursive nature of the formula, the power iteration method included in the Python package NetworkX^101^ was used with a convergence cutoff of 10^-6^. While EC’s can be calculated per eigen solution arising from eigendecomposition of the adjacency matrix, only the solution corresponding to the highest eigenvalue is considered^68,109^. This is in accordance with the Perron-Frobenius theorem that attributes uniqueness to the leading eigenvalue and its corresponding eigenvector calculated for a non-negative real square matrix (such as the adjacency matrix *A*) ^68,109-111^.

## Author contributions

CLW and LKM designed the studies and visualized the project. CLW performed the simulations and network maps. CLW, APK and LKM analyzed the data. CLW and LKM collated all data and wrote the manuscript. CLW, APK and LKM reviewed and edited the manuscript. Both authors approve the submitted version.

