## Supplementary Materials for "Allostery in Protein Tyrosine Phosphatases is Enabled by Divergent Dynamics"

#### Supplementary Figures

**Not for use without permission: Anyone who wishes to share, reuse, remix, or adapt this material must obtain permission from the corresponding author.**

**Supplementary Figure 1.** Structural and sequence comparison of the three enzymes under study. (A) Superimposition of the three starting structures for TbPTP1 (PDB ID: 3M4U in red), PTP1B ( PDB ID: 1PTV in gray) and YopH (PDB ID: 2YDU in pink). (B) Structure-based sequence alignment of the three proteins. The 10 canonical PTP motifs are highlighted in blue.

**Supplementary Figure 2.** Evolutionary analysis of the PTP domain (A) Phylogeny and evolutionary relationships of all solved PTP domain structures. Entries marked with a red dot are used in this study. (B) Conservation of physico-chemical properties, including charge, hydrophobicity, etc over the ~42,000 sequences from the PFAM database mapped onto the PTP domain. Lowest conservation (green) to highest conservation (purple) is shown on a color scale. (C) Percent consensus for residue type over each position for the ~42,000 sequences from the PFAM database mapped onto the PTP domain. Color scale is used to show the minimum at 0% (in green) to 50% (in white) to a maximum of 100% (in purple).

**Supplementary Figure 3.** Reaction mechanism of hydrolysis of substrate phosphotyrosines at the PTP active site. (A) Conserved catalytic mechanism for PTP enzymatic activity. Substrate atoms and bonds in dark red, enzyme atoms and bonds are in gray. (B) Kinetic parameters for TbPTP1, PTP1B and YopH as obtained from literature.

**Supplementary Figure 4.**Time evolution of structural properties of TbPTP1 as seen in the Molecular Dynamics Simulation runs. Root-mean square deviation (RMSD), Radius of gyration (Rg), and Solvent Accessible Surface Area (SASA) are shown from top to bottom. Apo and pY bound TbPTP1 are shown separately. Data for all four runs is shown for each parameter.

**Supplementary Figure 5.** Normalized probability density of structural factors of Root-mean square deviation (RMSD), Radius of gyration (Rg), and Solvent Accessible Surface Area (SASA) for the equilibrated trajectories of TbPTP1 and TbPTP1 in complex with pY. Data of all four trajectories is shown for each case.

**Supplementary Figure 6.** Time evolution of structural properties of PTP1B as seen the Molecular Dynamics Simulation runs. Root-mean square deviation (RMSD), Radius of gyration (Rg), and Solvent Accessible Surface Area (SASA) are shown from top to bottom. Apo and pY bound PTP1B are shown separately. Data for all four runs is shown for each parameter.

**Supplementary Figure 7.** Normalized probability density of structural factors of Root-mean square deviation (RMSD), Radius of gyration (Rg), and Solvent Accessible Surface Area (SASA) for the equilibrated trajectories of PTP1B and PTP1B in complex with pY. Data of all four trajectories is shown for each case.

**Supplementary Figure 8.** Time evolution of structural properties of YopH as seen the Molecular Dynamics Simulation runs. Root-mean square deviation (RMSD), Radius of gyration (Rg), and Solvent Accessible Surface Area (SASA) are shown from top to bottom. Apo and pY bound YopH are shown separately. Data for all four runs is shown for each parameter.

**Supplementary Figure 9.** Normalized probability density of structural factors of Root-mean square deviation (RMSD), Radius of gyration (Rg), and Solvent Accessible Surface Area (SASA) for the equilibrated trajectories of YopH and YopH in complex with pY. Data of all four trajectories is shown for each case.

**Supplementary Figure 10.** Residue wise Root Mean Square Fluctuations (RMSF) as seen during the simulation for the three proteins: TbPTP1 (top), PTP1B (middle), and YopH (bottom). Both Apo and pY-bound systems are shown right). Catalytic motifs are highlighted: pY-loop (yellow), E-loop (red), WPD-loop (orange), P-loop (green), and Q-loop (blue).

**Supplementary Figure 11.** Essential dynamics of the Apo and pY-bound forms of TbPTP1, PTP1B and YopH as seen by PCA analysis. Contribution of each eigenvector/mode to the total variance of simulations is shown for the total set of eigenvectors (top) and the first 10 eigenvectors (bottom).

**Supplementary Figure 12.** RMSF plots for the first 10 eigenvectors of PCA for TbPTP1 (top), PTP1B (middle), and YopH (bottom) systems in the Apo (left) and pY bound (right) forms. Catalytic motifs are highlighted: pY-loop (yellow), E-loop (red), WPD-loop (orange), P-loop (green), and Q-loop (blue).

**Supplementary Figure 13.** First five PCA eigenvectors mapped onto the structure of TbPTP1. TbPTP1-apo modes (left) and TbPTP1+pY modes (right) are shown. The initial frame of each mode is colored orange and final frame colored pink.

**Supplementary Figure 14.** First five PCA eigenvectors mapped onto the structure of PTP1B. PTP1B-apo modes (left) and PTP1B+pY modes (right) are shown. The initial frame of each mode is colored orange and final frame colored pink.

**Supplementary Figure 15.** First five PCA eigenvectors mapped onto the structure of YopH. YopH-apo modes (left) and YopH+pY modes (right) are shown. The initial frame of each mode is colored orange and final frame colored pink.

**Supplementary Figure 16.** PCA of Cα motions. (A) Overlay of 5 first individual modes of motion between the apo (left column) and pY-bound (right column) states. Motions are shown for all three enzymes, TbPTP1 (top), PTP1B (middle), and YopH (bottom). Initial frames for each individual mode are in orange, the final frame in pink. Arrows in cyan show motions of Cα atoms > 1.5Å. (B) Cosine similarity index for each pair of vectors for a given enzyme.

**Supplementary Figure 17.** Normalized probability density distributions for various distances seen in the active sites of TbPTP1 (top), PTP1B (middle), and YopH (bottom). All distances are measured between corresponding $C_\alpha$ backbone atoms. Distances shown are for the pY-loop – P-loop (first column; TbPTP1: Y51-C229; PTP1B: Y46-C215; YopH: F229-C403), the Q-loop – P-loop (second column; TbPTP1: Q275-C229; PTP1B: Q262-C215; YopH: Q446-C403), the pY-loop – WPD-loop (third column; TbPTP1: Y51-D199; PTP1B: Y46-D181; YopH: F229-D356), and the P-loop – WPD-loop (fourth TbPTP1: C229-D199; PTP1B: C215-D181; YopH: C403-D356). Data for of all four trajectories of both Apo and pY-bound states of the three proteins is shown.

**Supplementary Figure 18.** Normalized probability density distributions for the dihedral angles of the P-Loop arginine as seen in the Apo and pY-bound simulations of the TbPTP1, PTP1B and YopH. Plots for the Apo state (purple) and pY-bound state (orange) are shown for TbPTP1 (left; R235), PTP1B (middle, R221), and YopH (right, R409).

**Supplementary Figure 19.** Community Analysis and influential nodes. (A) Girvan-Newman communities of TbPTP1-apo (top) and TbPTP1+pY (bottom) mapped to the corresponding maximum density structure obtained from simulations (as shown in Figure 5B of the main text). The ribbon depiction is colored according to the community of the main chain node, which the corresponding sidechain node appearing as a ball-and-stick representation of the $C_\beta$ atom and colored according to its own community. (B) Nodes with the highest Eigen centrality (from Figure 7 in the main text) mapped to the community structure of TbPTP1-apo (top) and TbPTP1+pY (bottom). Also see Supplementary Figures 22 & 23.

**Supplementary Figure 20.** Community Analysis and influential nodes. (A) Girvan-Newman communities of PTP1B-apo (top) and PTP1B+pY (bottom) mapped to the corresponding maximum density structure obtained from simulations (as shown in Figure 5B of the main text). The ribbon depiction is colored according to the community of the main chain node, which the corresponding sidechain node appearing as a ball-and-stick representation of the $C_\beta$ atom and colored according to its own community. (B) Nodes with the highest Eigen centrality (from Figure 7 in the main text) mapped to the community structure of PTP1B-apo (top) and PTP1B+pY (bottom). Also see Supplementary Figures 24 & 25.

**Supplementary Figure 21.** Community Analysis and influential nodes. (A) Girvan-Newman communities of YopH-apo (top) and YopH+pY (bottom) mapped to the corresponding maximum density structure obtained from simulations (as shown in Figure 5B of the main text). The ribbon depiction is colored according to the community of the main chain node, which the corresponding sidechain node appearing as a ball-and-stick representation of the $C_{\beta}$ atom and colored according to its own community. (B) Nodes with the highest Eigen centrality (from Figure 7 in the main text) mapped to the community structure of YopH-apo (top) and YopH+pY (bottom). Also see Supplementary Figures 26 & 27.

**Supplementary Figure 22.** Whole network visualization for TbPTP1-apo. (a) ForceAtlas2 distribution of nodes colored by Girvan-Newman community assignment. The size of the nodes corresponds to the eigencentrality value of that node. Nodes with eigencentrality < 0.001 are set to size 0.001 for ease of viewing. Community 2 is excluded to better fit the image. (B) Zoomed-in cluster of high-eigencentrality nodes from (A).

**Supplementary Figure 23.** Whole network visualization for TbPTP1+pY. (a) ForceAtlas2 distribution of nodes colored by Girvan-Newman community assignment. The size of the nodes corresponds to the eigencentrality value of that node. Nodes with eigencentrality < 0.001 are set to size 0.001 for ease of viewing. Community 2 is excluded to better fit the image. (B) Zoomed-in cluster of high-eigencentrality nodes from (A).

**Supplementary Figure 24.** Whole network visualization for PTP1B-apo. (a) ForceAtlas2 distribution of nodes colored by Girvan-Newman community assignment. The size of the nodes corresponds to the eigencentrality value of that node. Nodes with eigencentrality < 0.001 are set to size 0.001 for ease of viewing. (B) Zoomed-in cluster of high-eigencentrality nodes from (A).

**Supplementary Figure 25.** Whole network visualization for PTP1B+pY. (a) ForceAtlas2 distribution of nodes colored by Girvan-Newman community assignment. The size of the nodes corresponds to the eigencentrality value of that node. Nodes with eigencentrality < 0.001 are set to size 0.001 for ease of viewing. (B) Zoomed-in clusters of high-eigencentrality nodes from (A).

**Supplementary Figure 26.** Whole network visualization for YopH-apo. (a) ForceAtlas2 distribution of nodes colored by Girvan-Newman community assignment. The size of the nodes corresponds to the eigencentrality value of that node. Nodes with eigencentrality < 0.001 are set to size 0.001 for ease of viewing. Community 1 is excluded to better fit the image. (B) Zoomed-in cluster of high-eigencentrality nodes from (A).

**Supplementary Figure 27.** Whole network visualization for YopH+pY. (a) ForceAtlas2 distribution of nodes colored by Girvan-Newman community assignment. The size of the nodes corresponds to the eigencentrality value of that node. Nodes with eigencentrality < 0.001 are set to size 0.001 for ease of viewing. Community 1 is excluded to better fit the image. (B) Zoomed-in cluster of high-eigencentrality nodes from (A).

#### Supplementary Figure 1

**A**

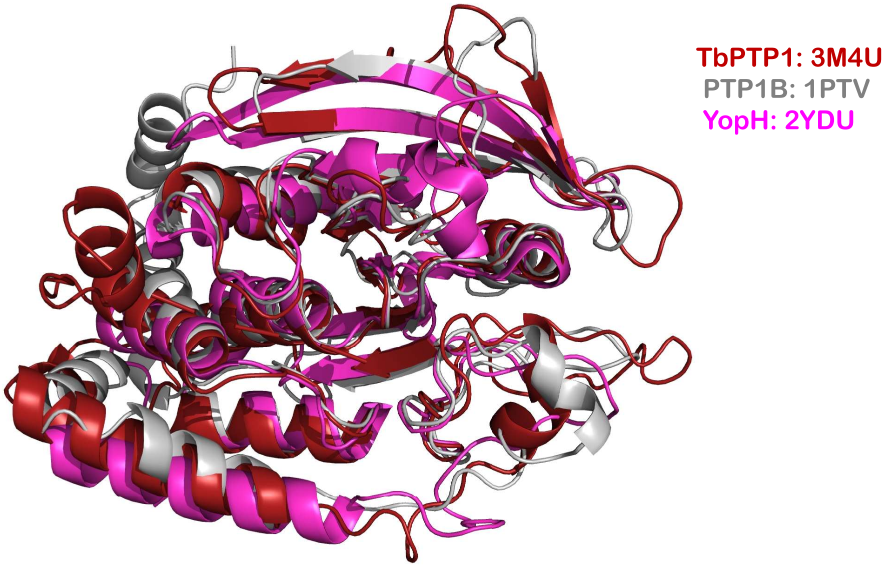


**B**

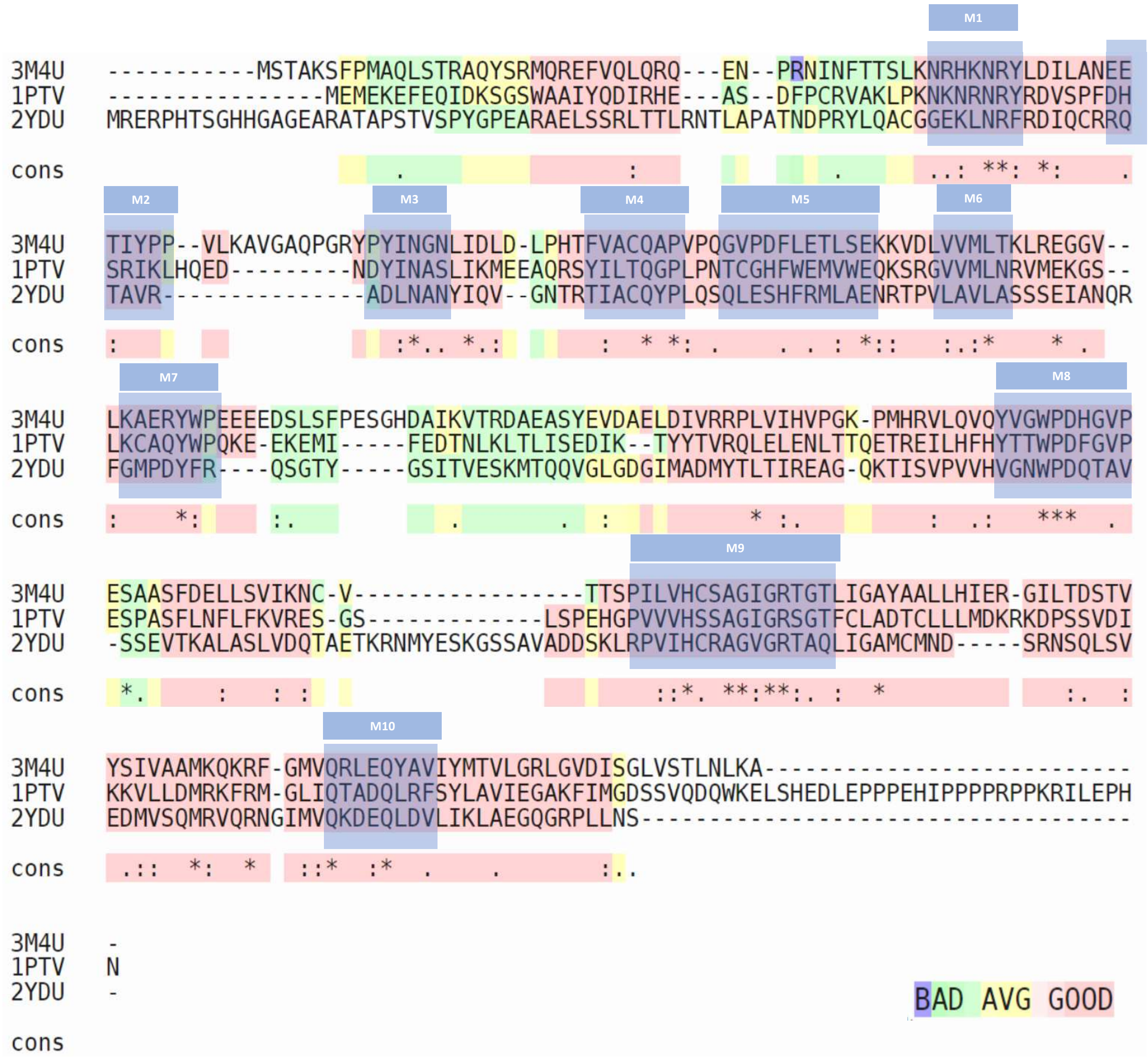

#### Supplementary Figure 2

A

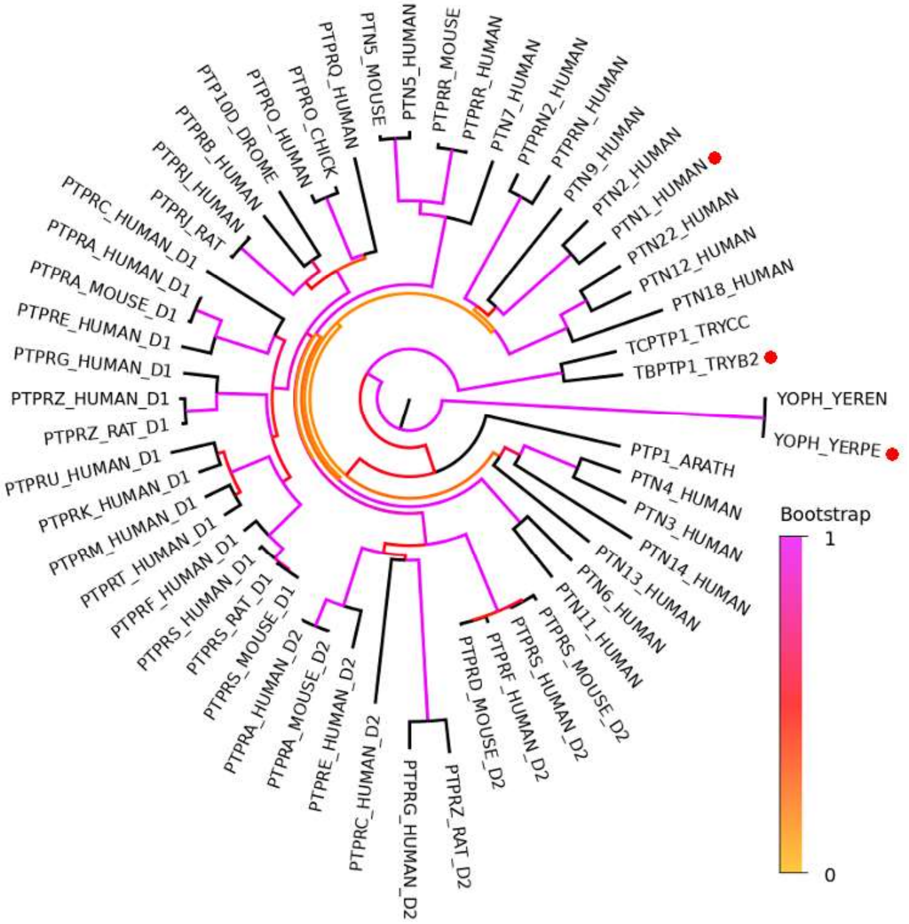


B

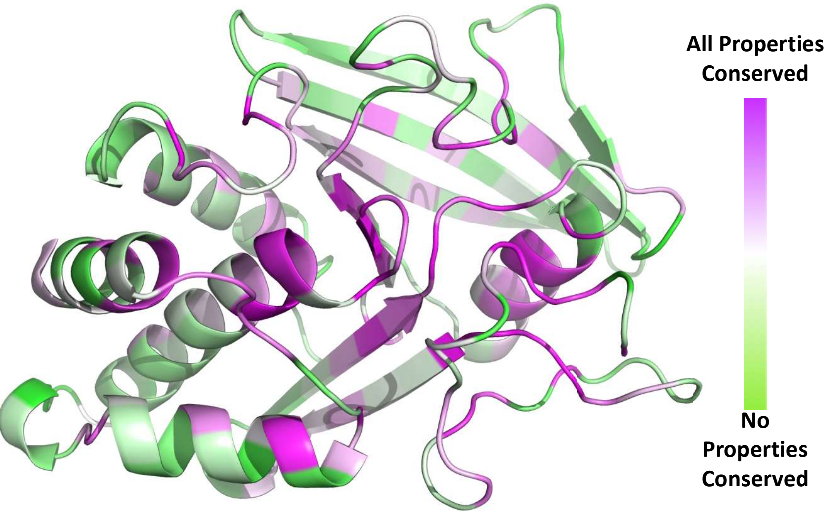


C

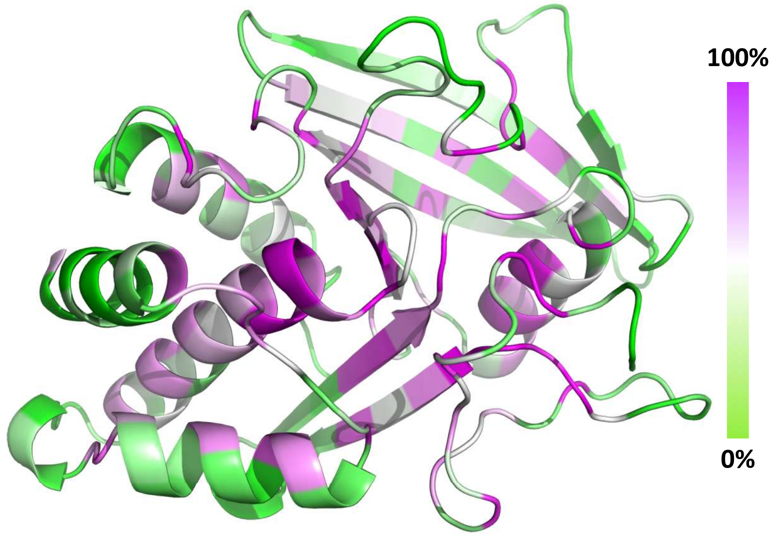

#### Supplementary Figure 3

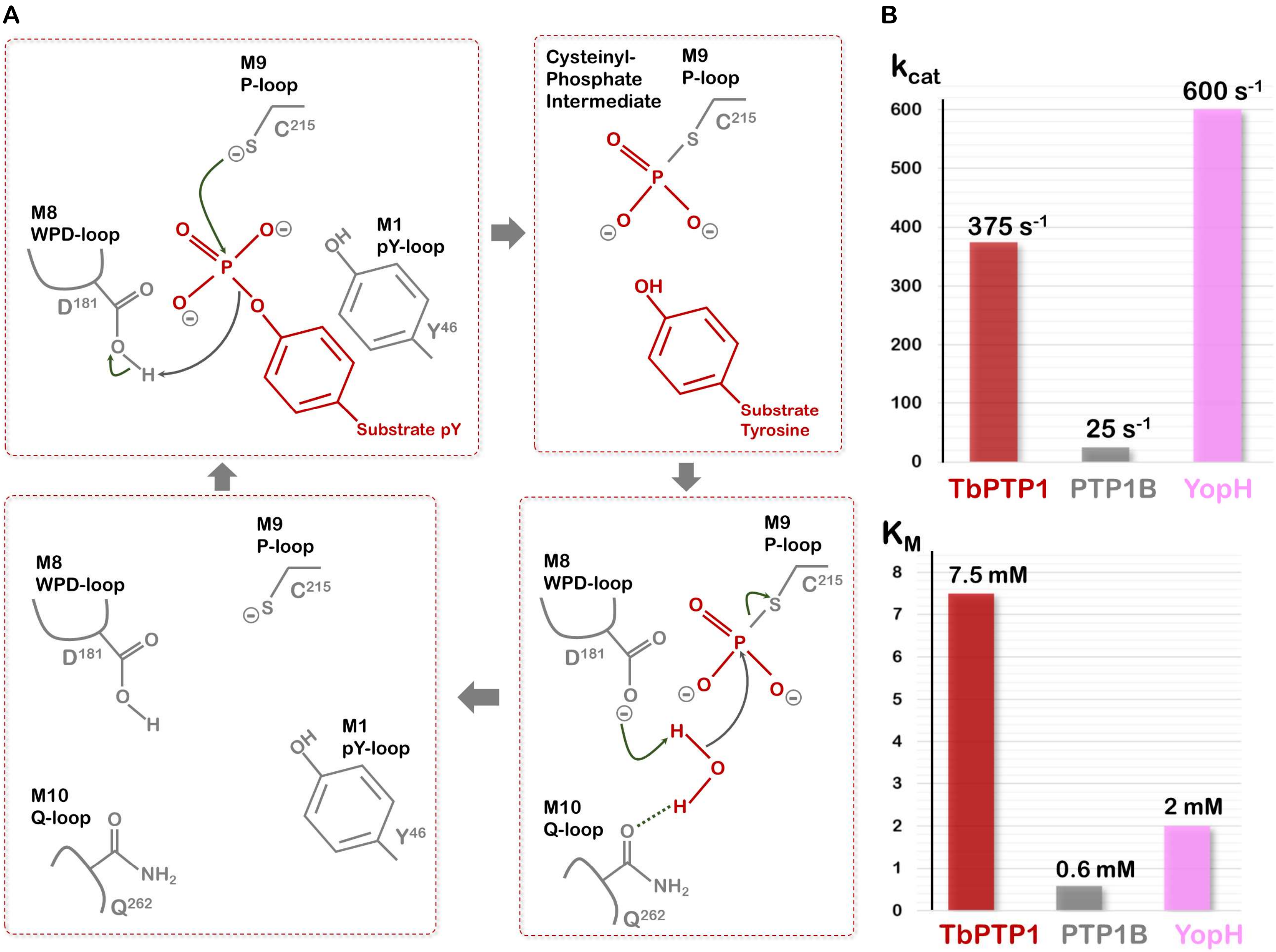

#### Supplementary Figure 4

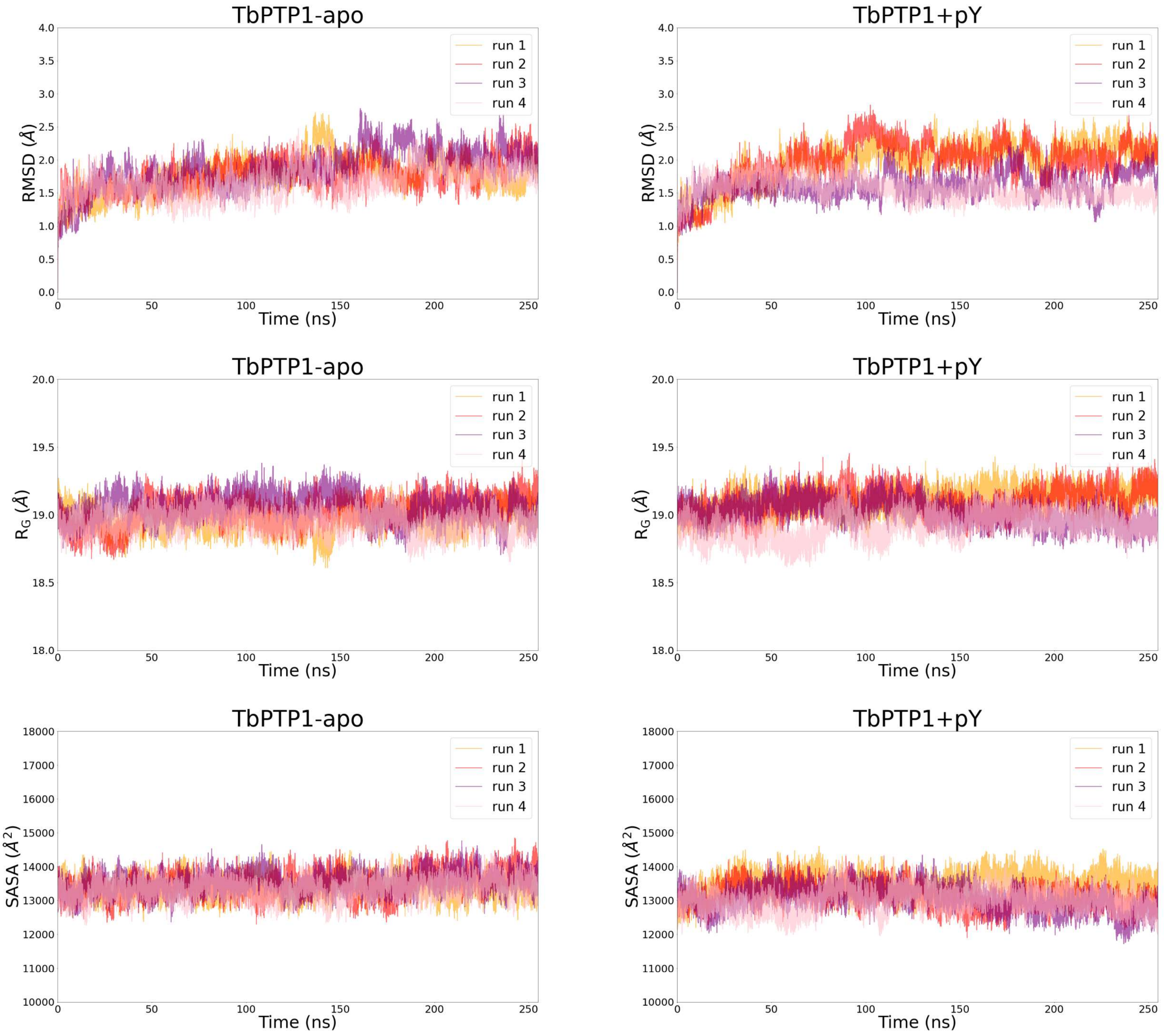

#### Supplementary Figure 5

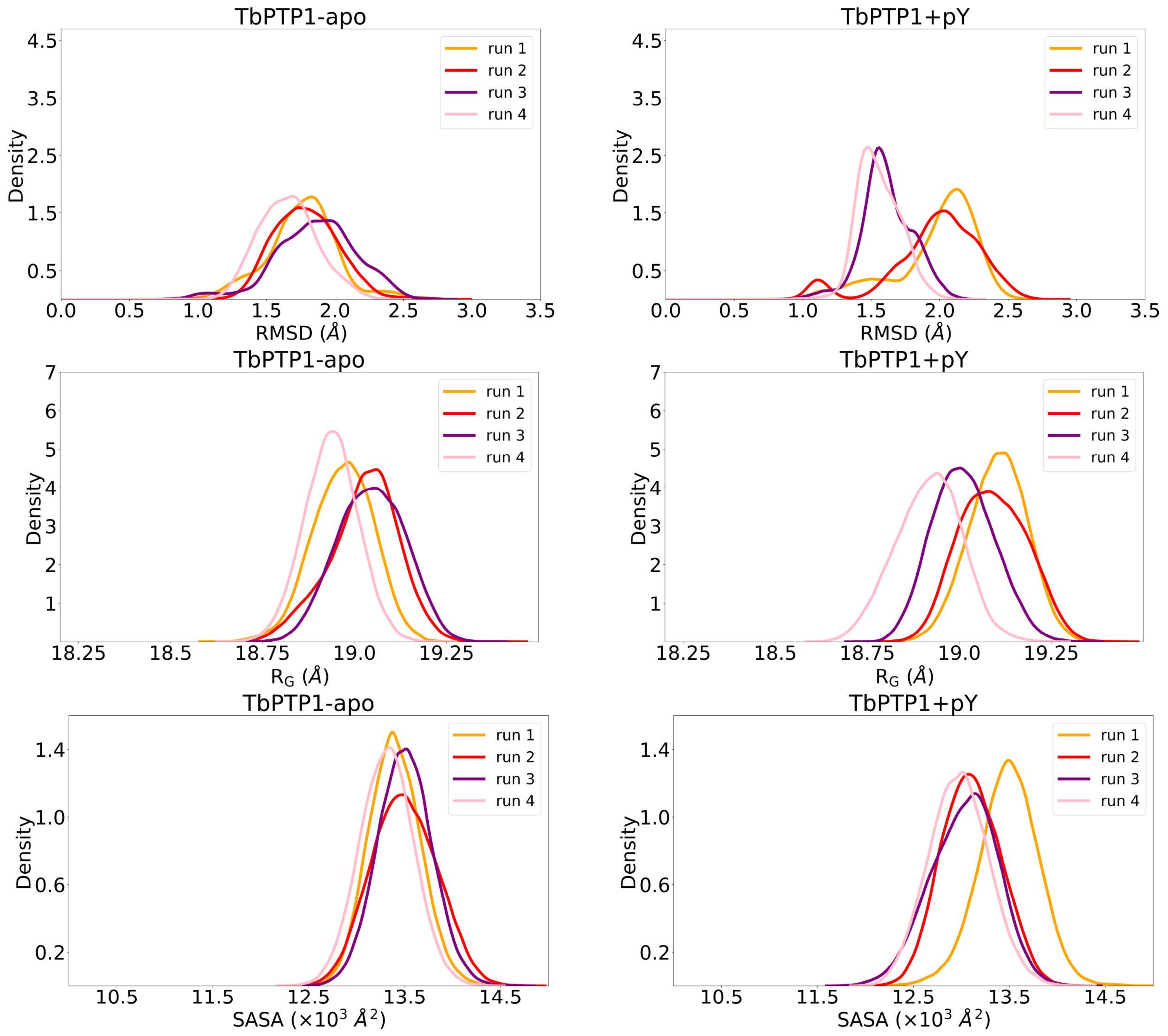

### Supplementary Figure 6

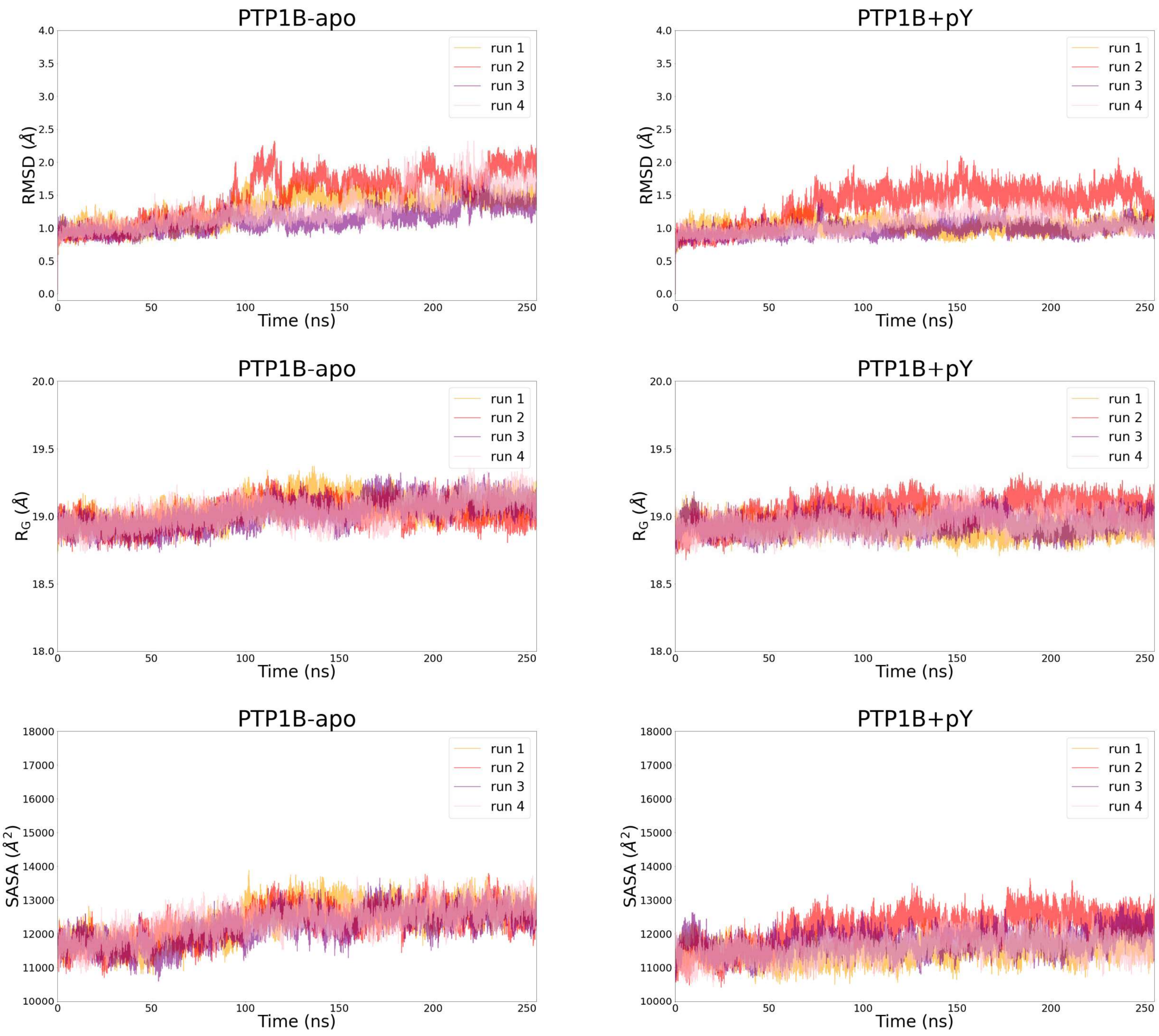

**Supplementary Figure 7**

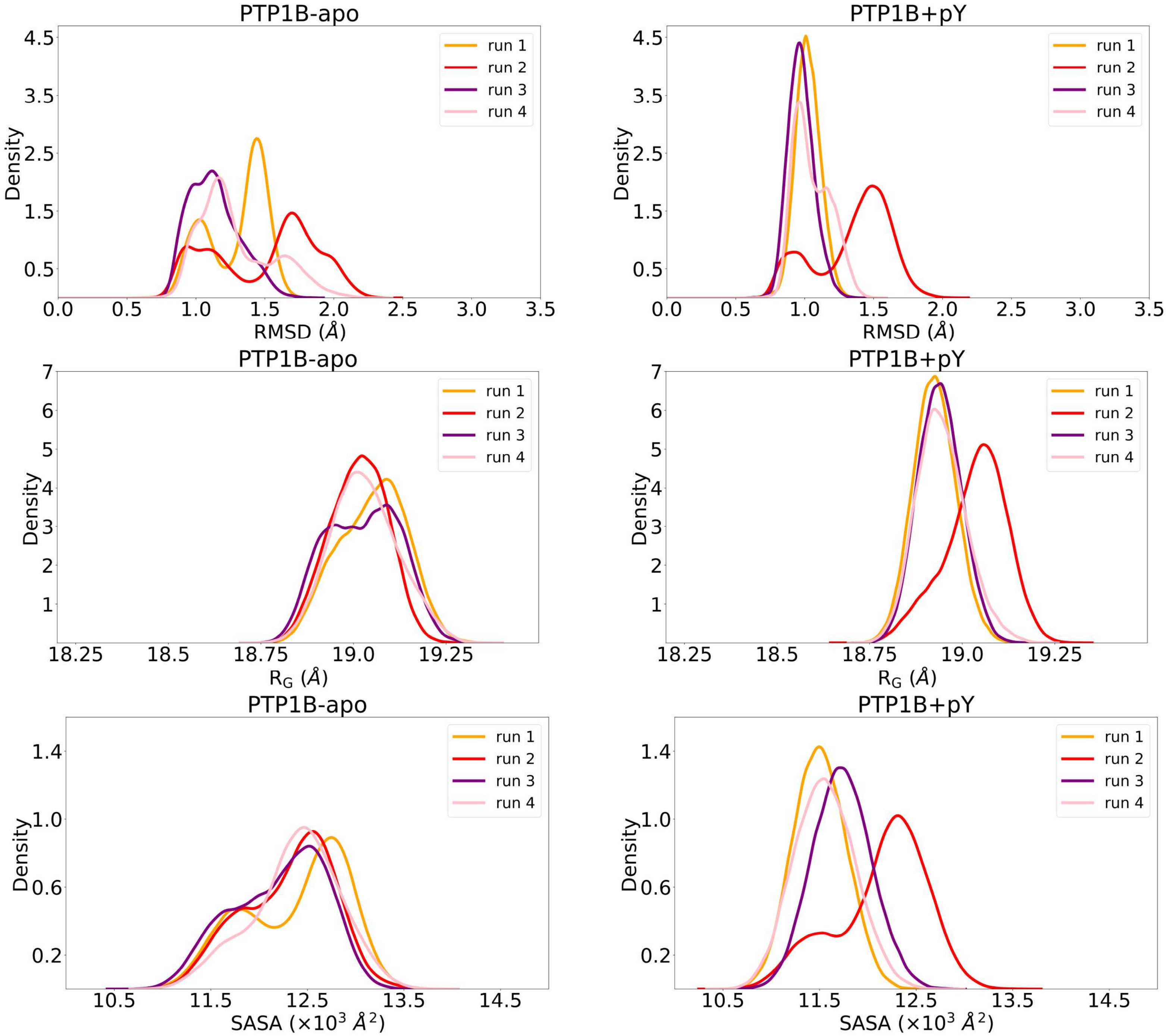

#### Supplementary Figure 8

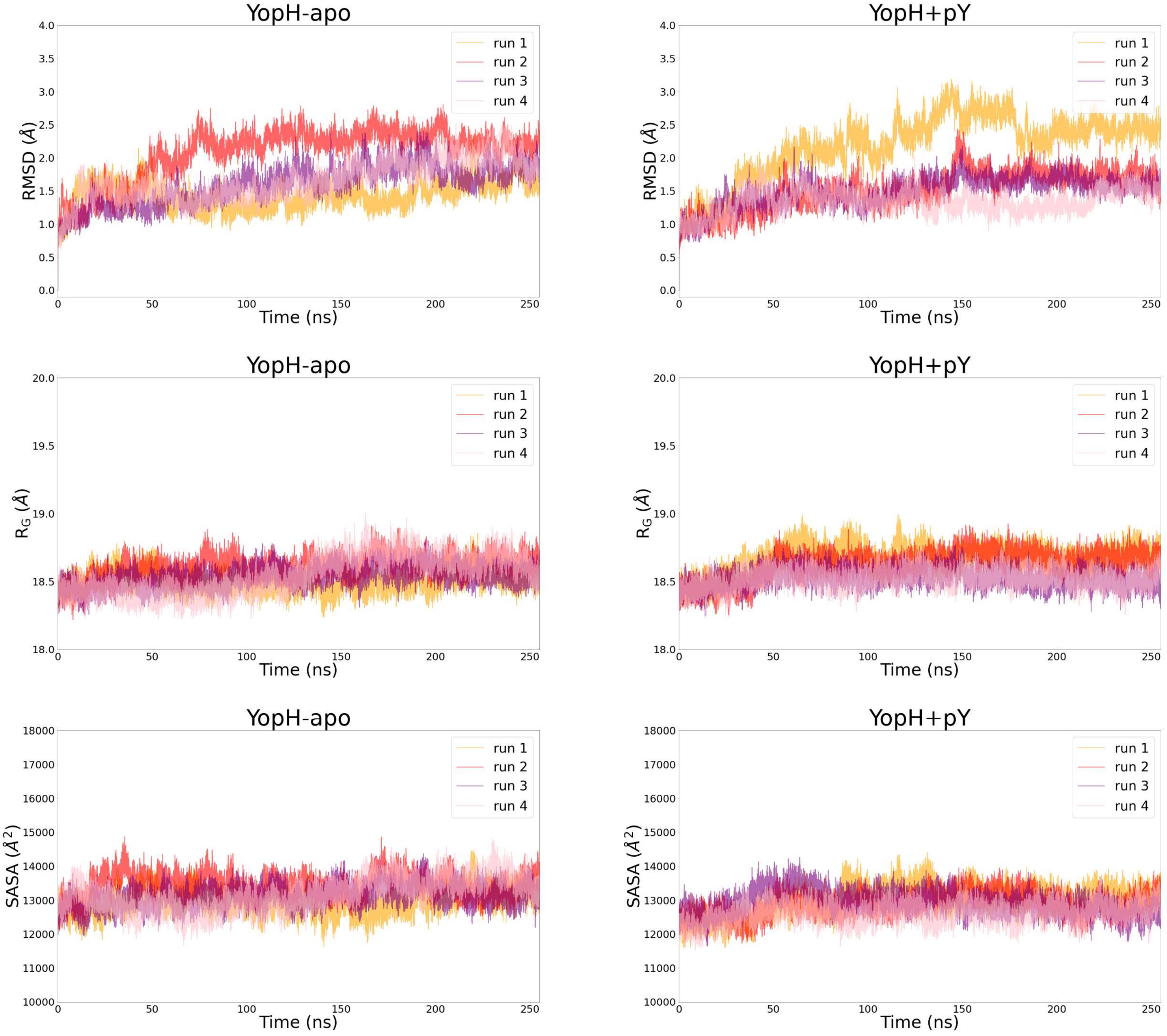

#### Supplementary Figure 9

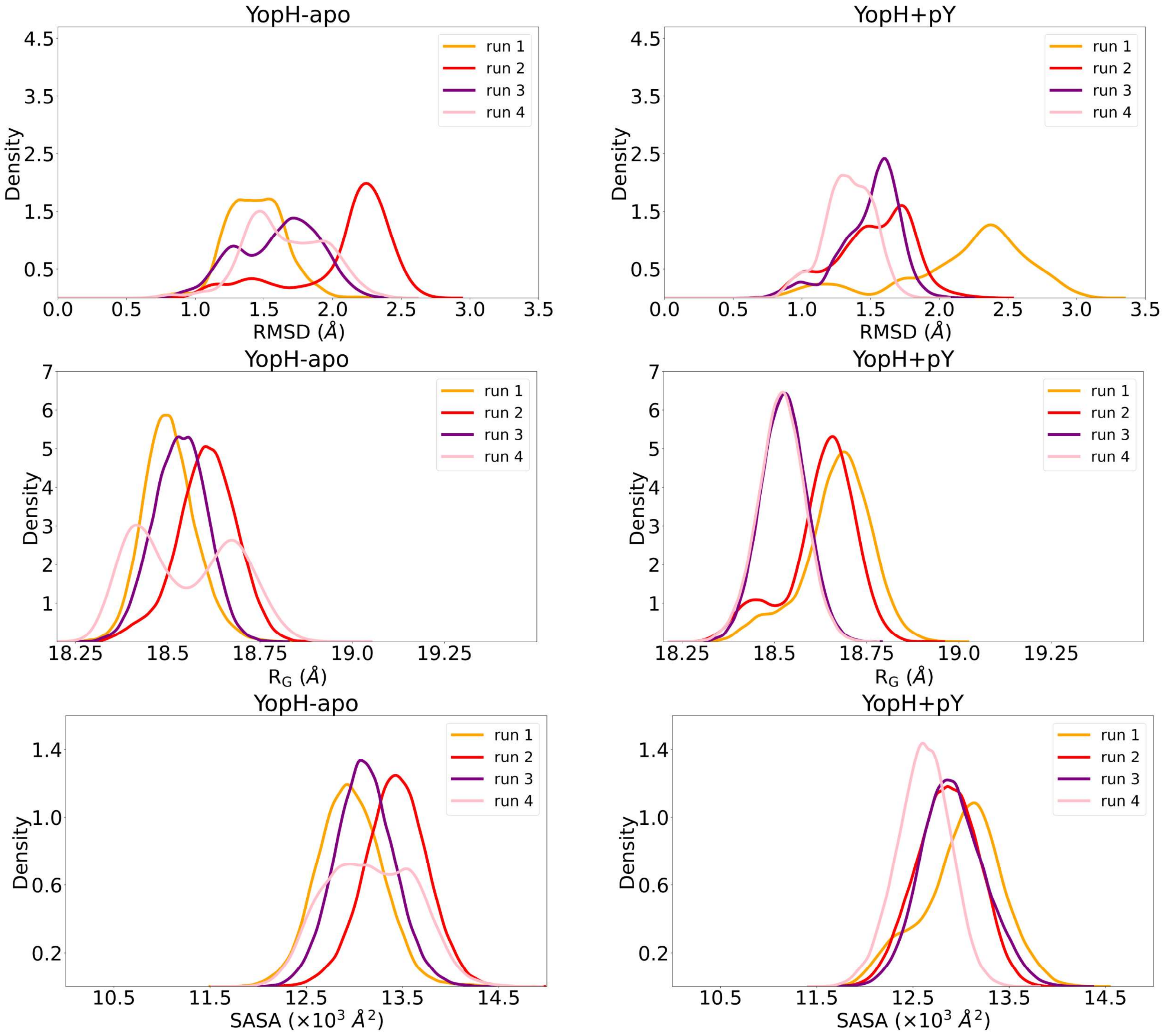

### Supplementary Figure 10

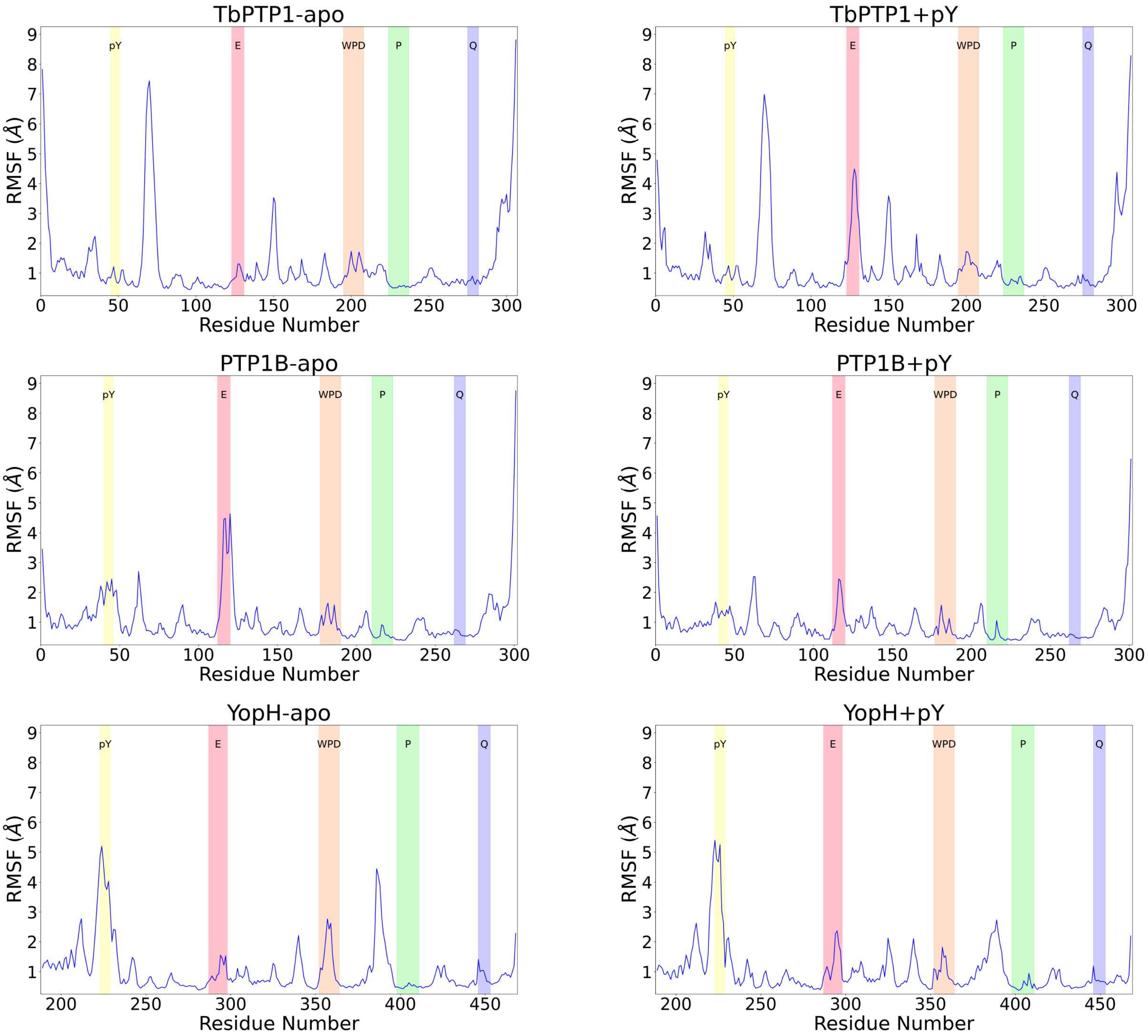

**Supplementary Figure 11**

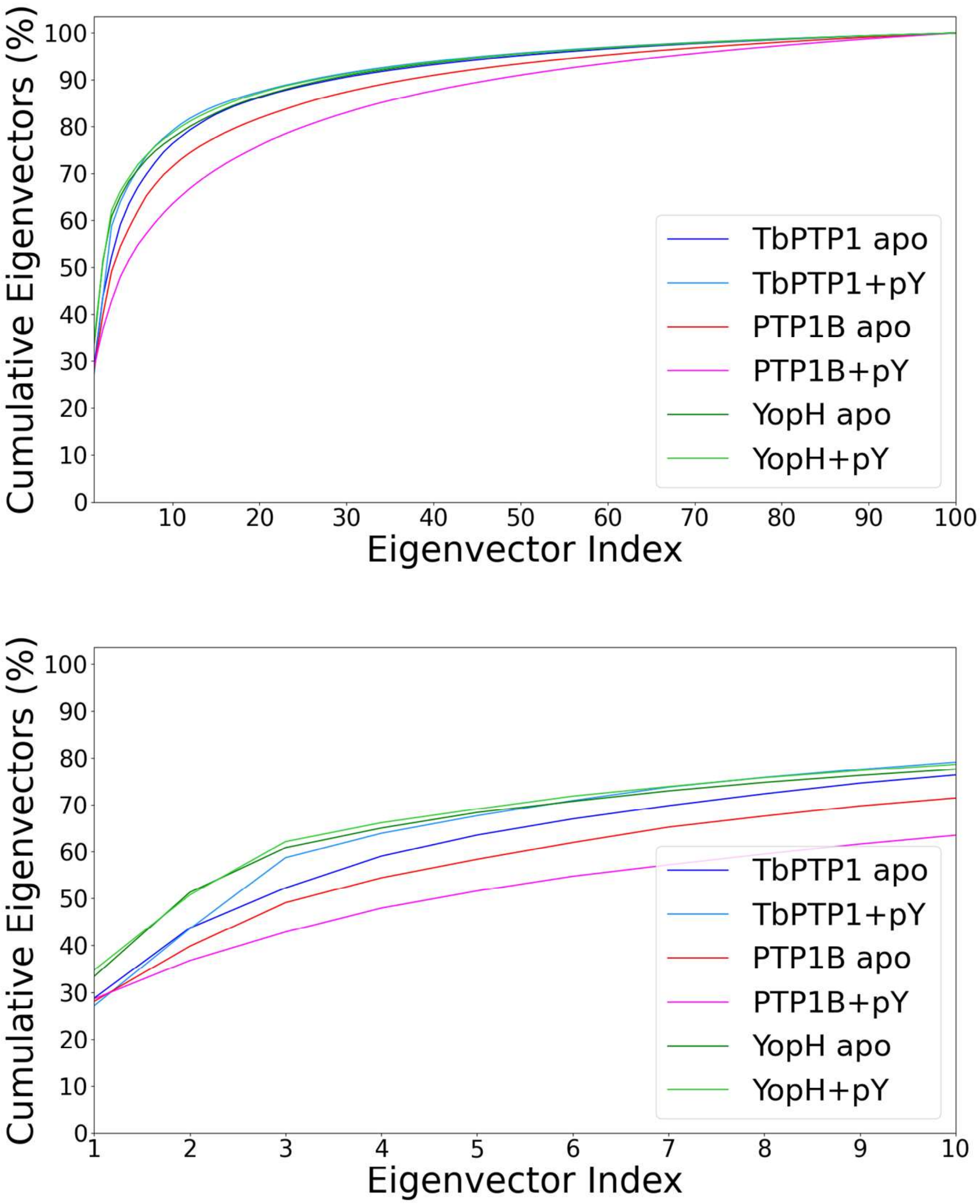

### Supplementary Figure 12

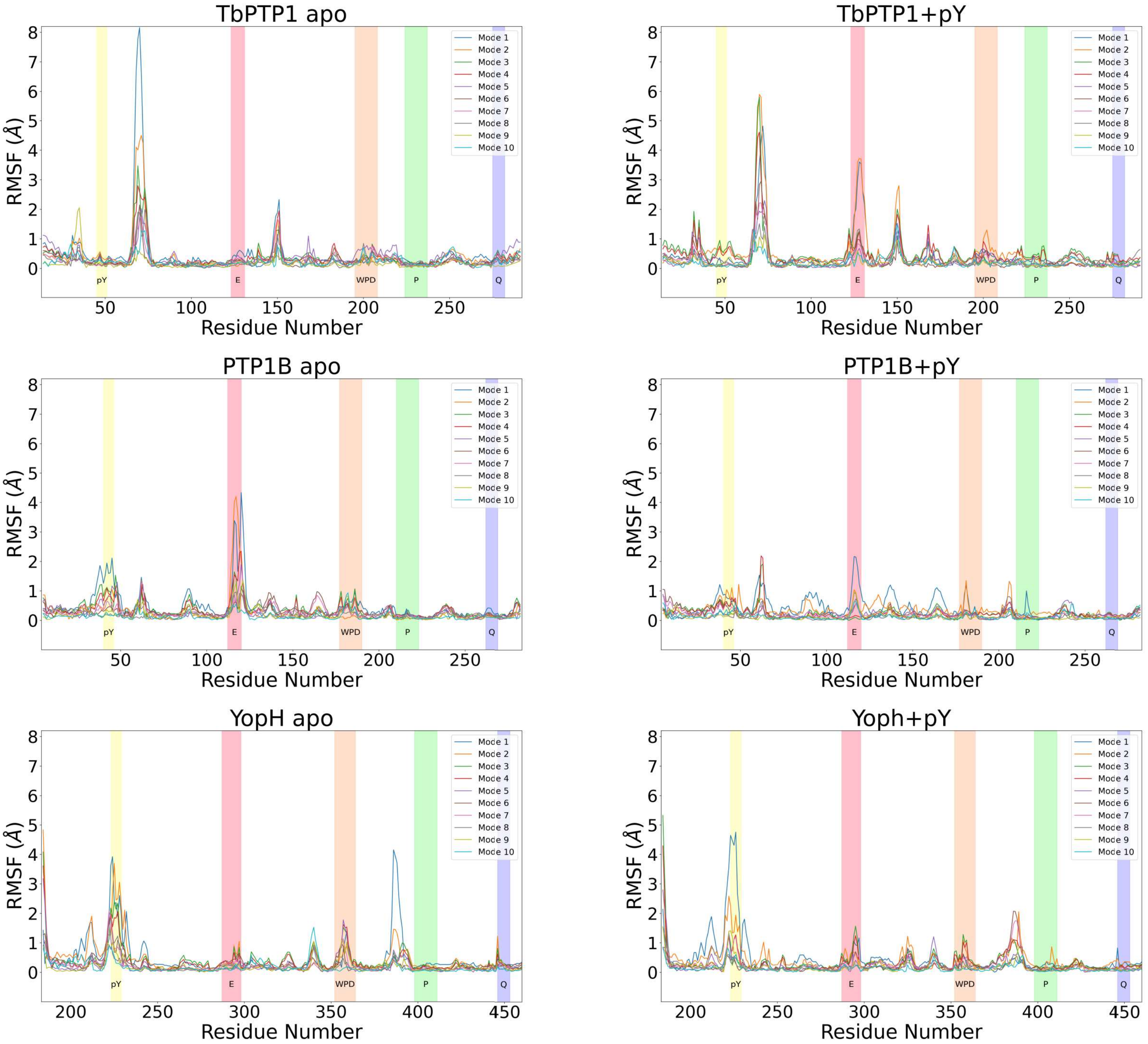

#### Supplementary Figure 13

Mode — TbPTP1 Apo — TbPTP1 + pY

1

2

3

4

5

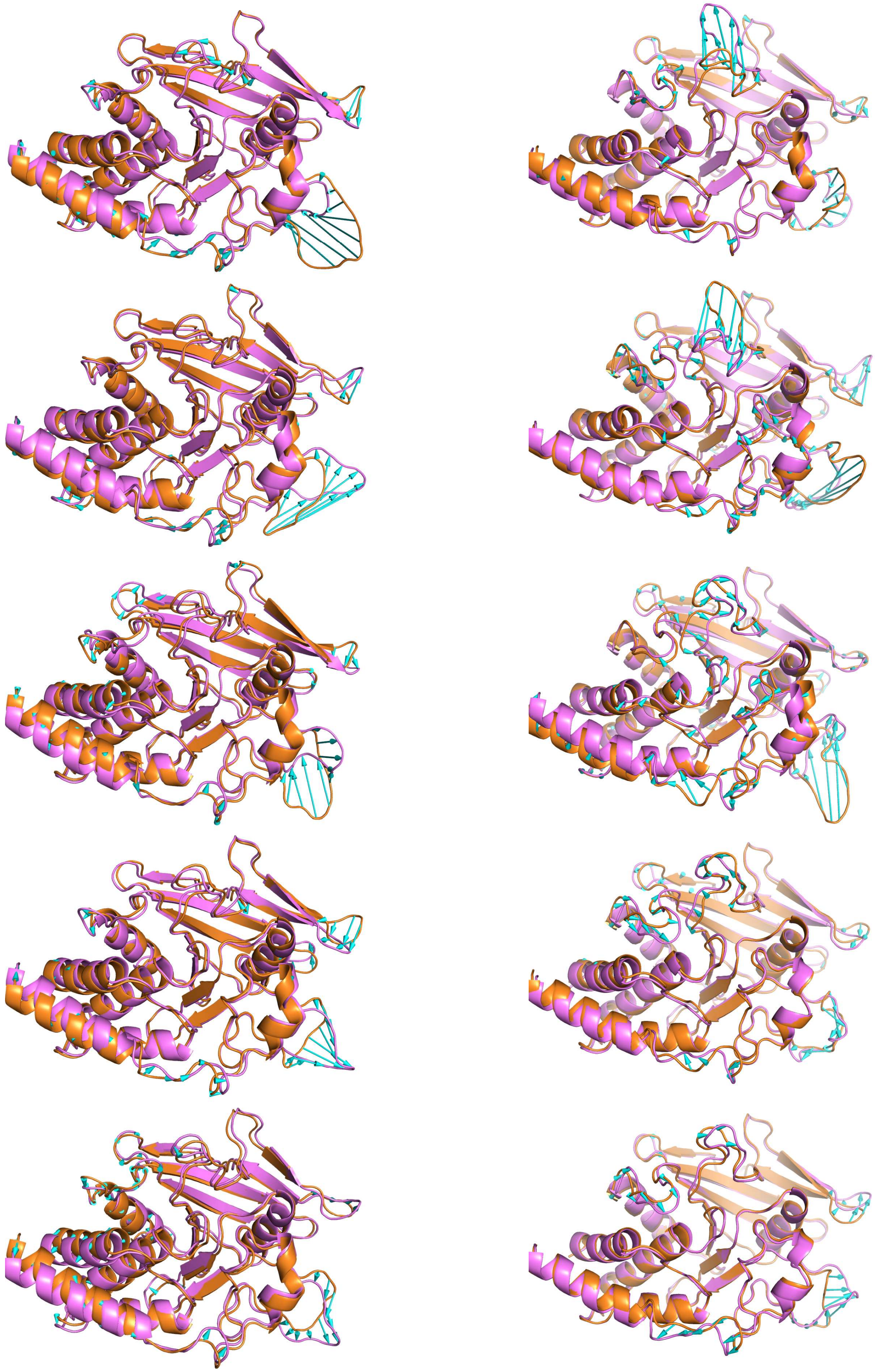

Supplementary Figure 14

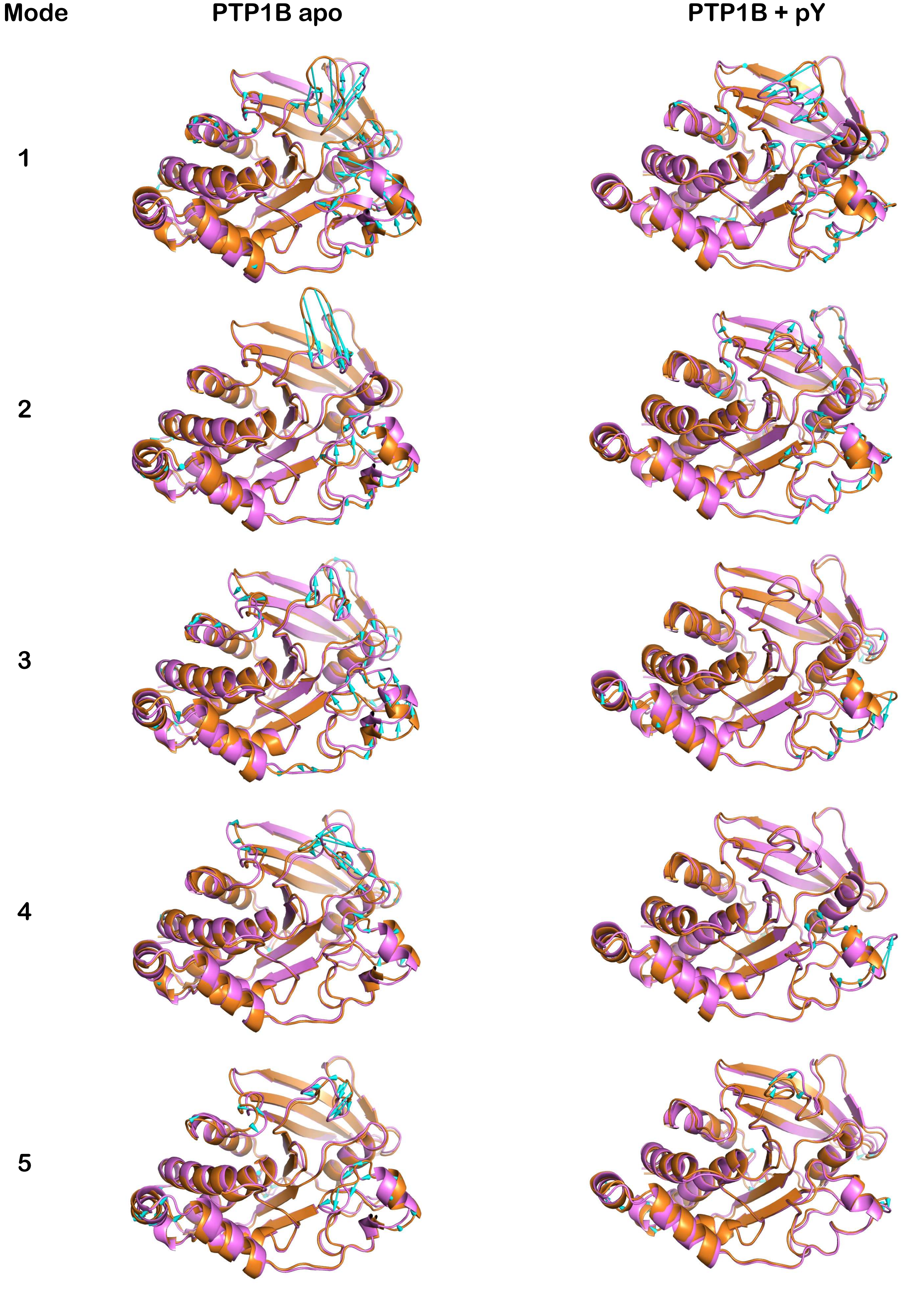

Supplementary Figure 15

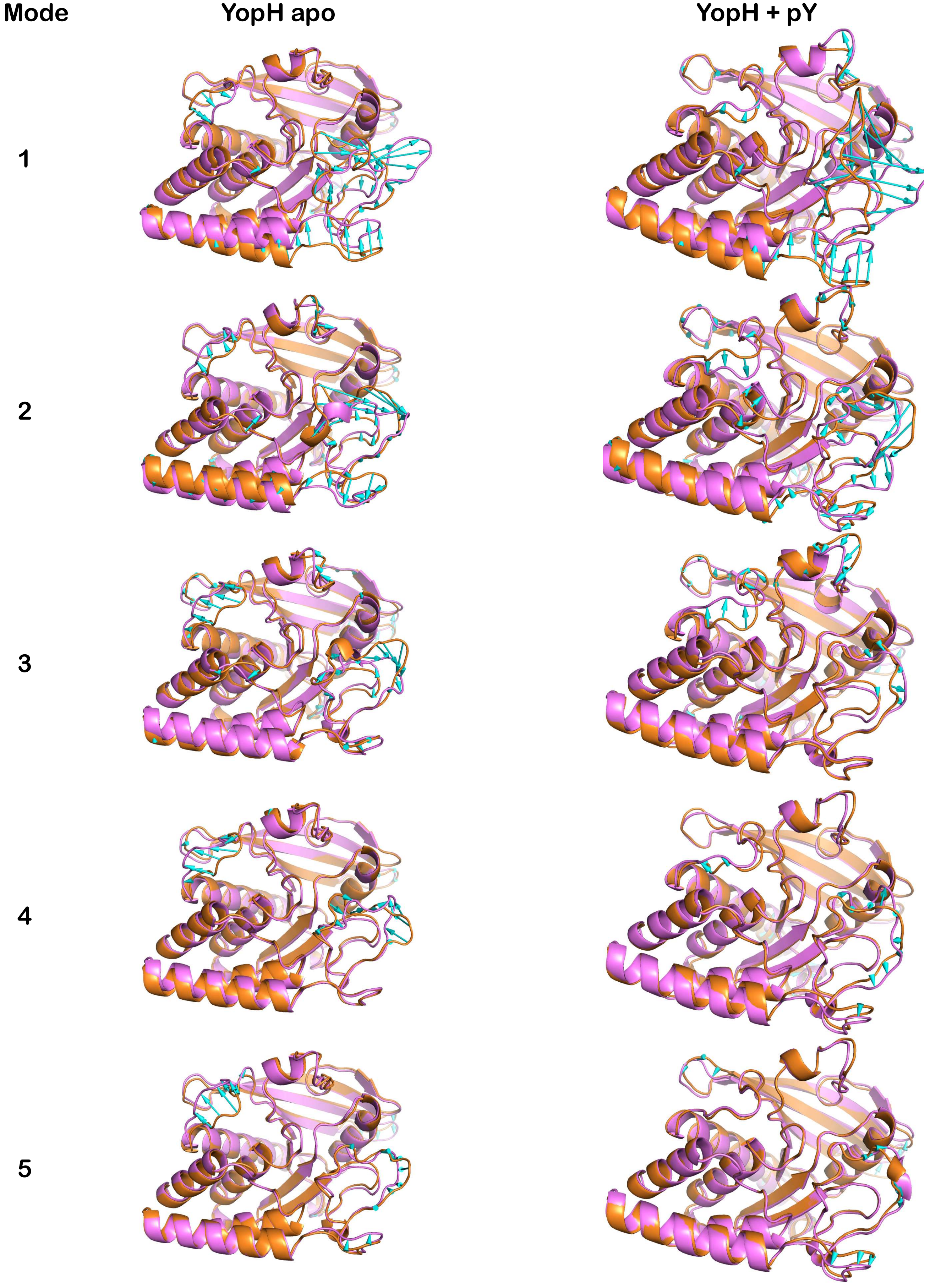

#### Supplementary Figure 16

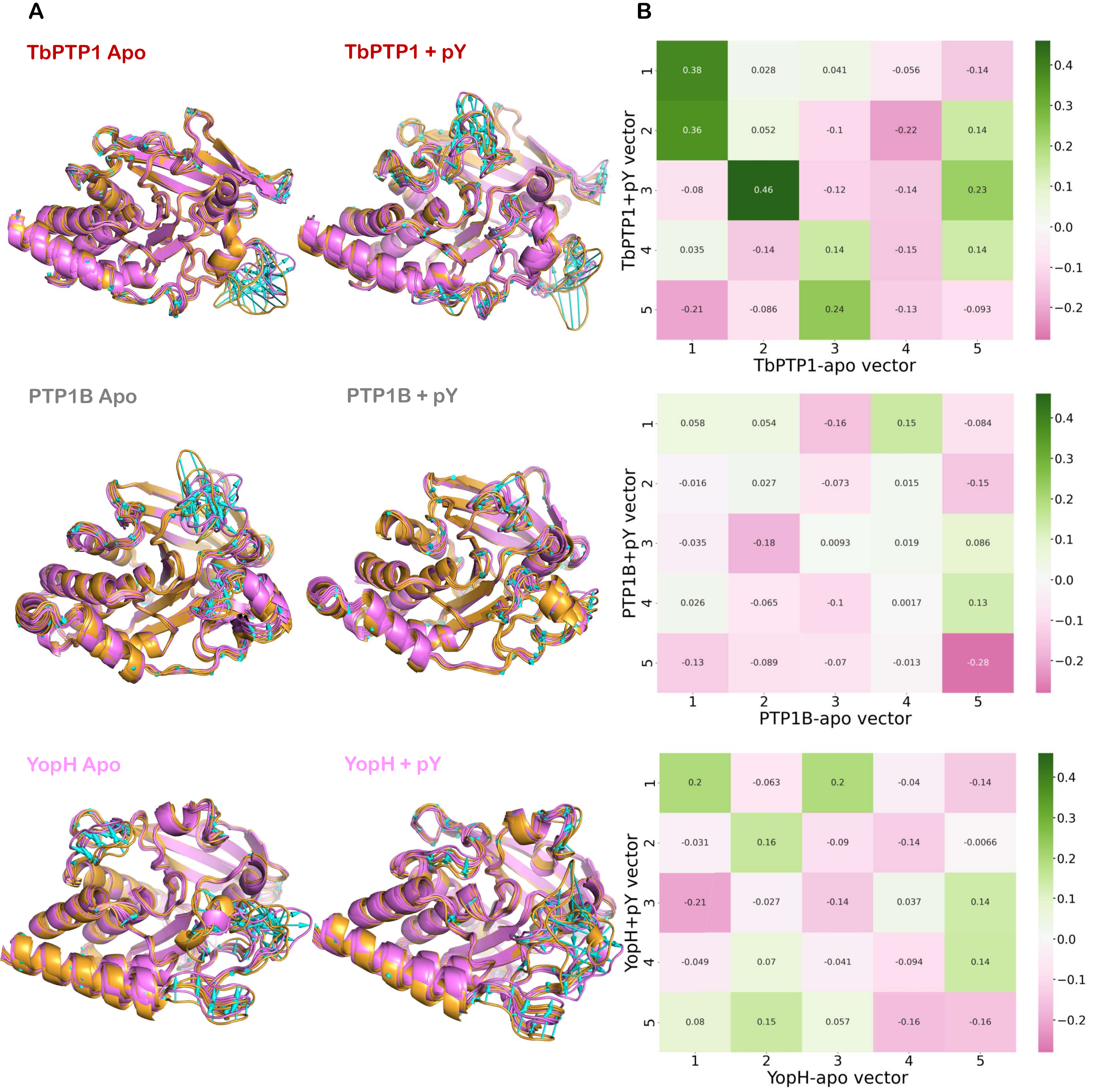

### Supplementary Figure 17

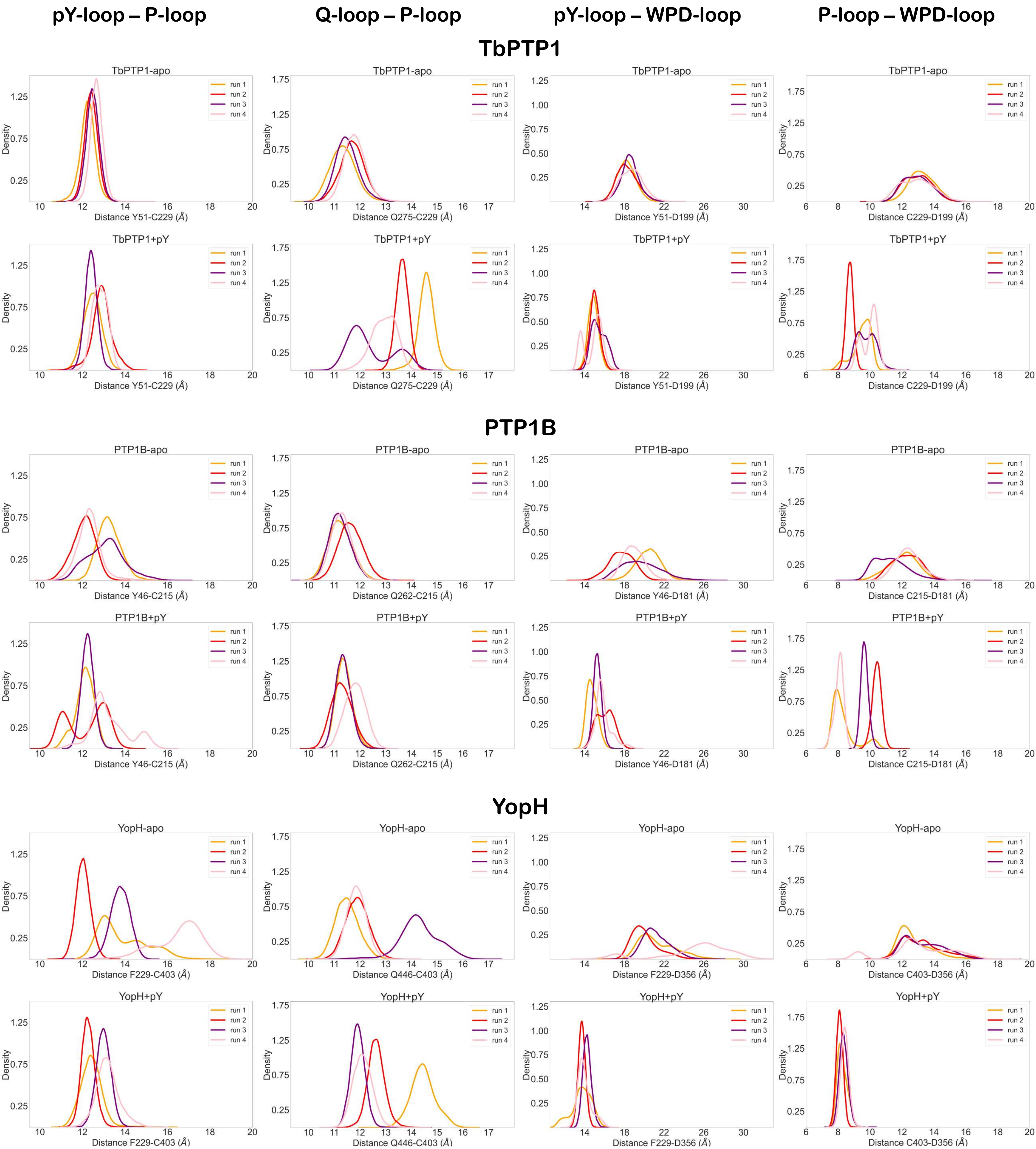

#### Supplementary Figure 18

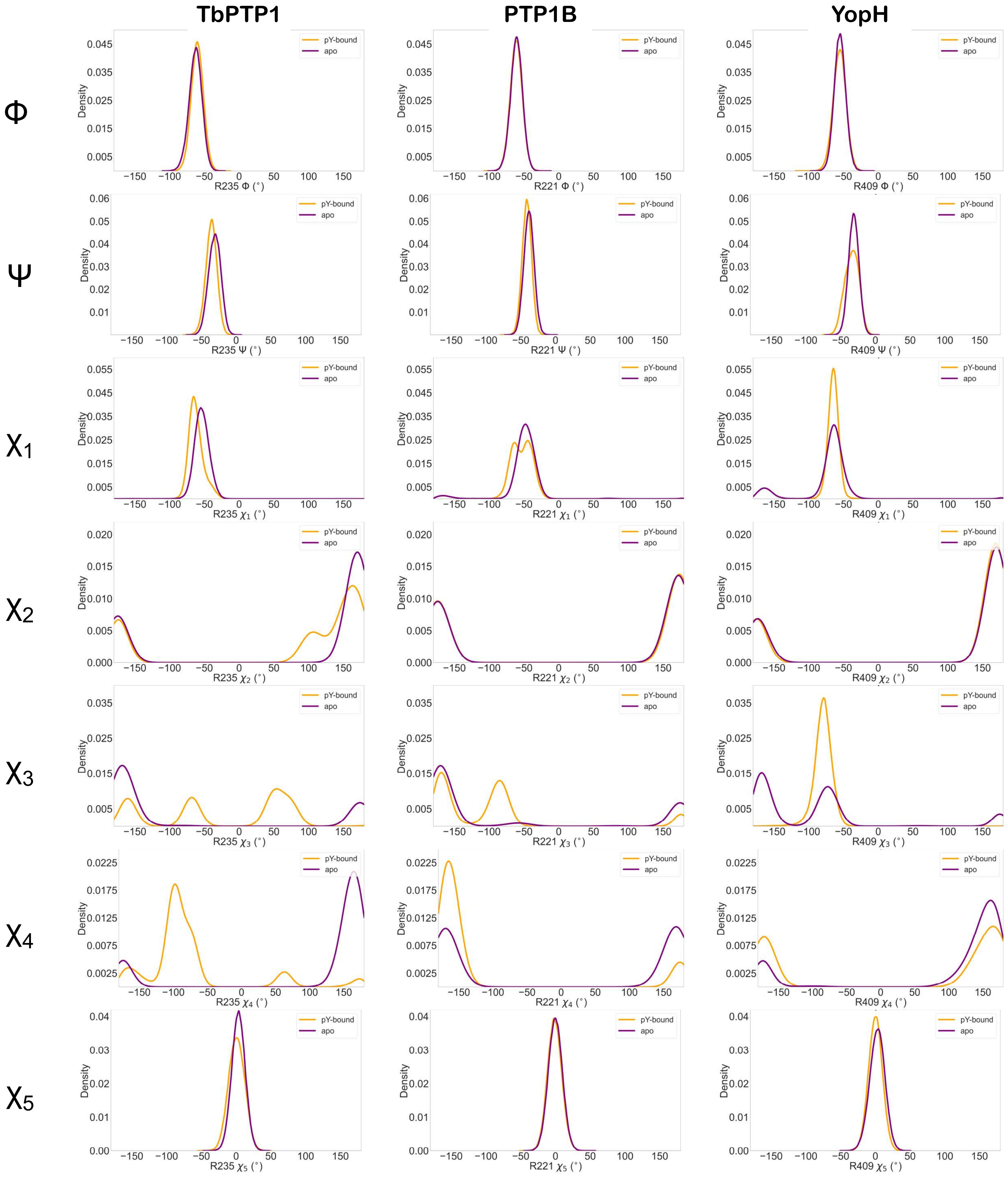

#### Supplementary Figure 19

**TbPTP1 apo**

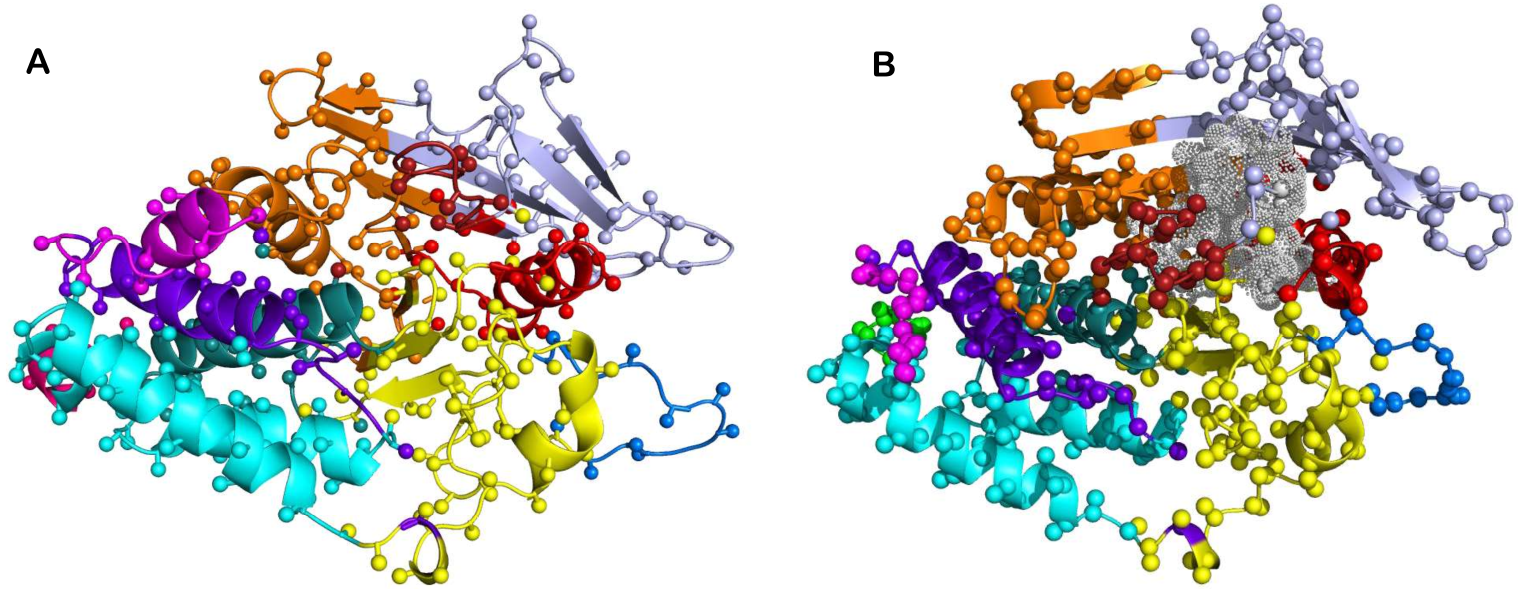


**TbPTP1 + pY**

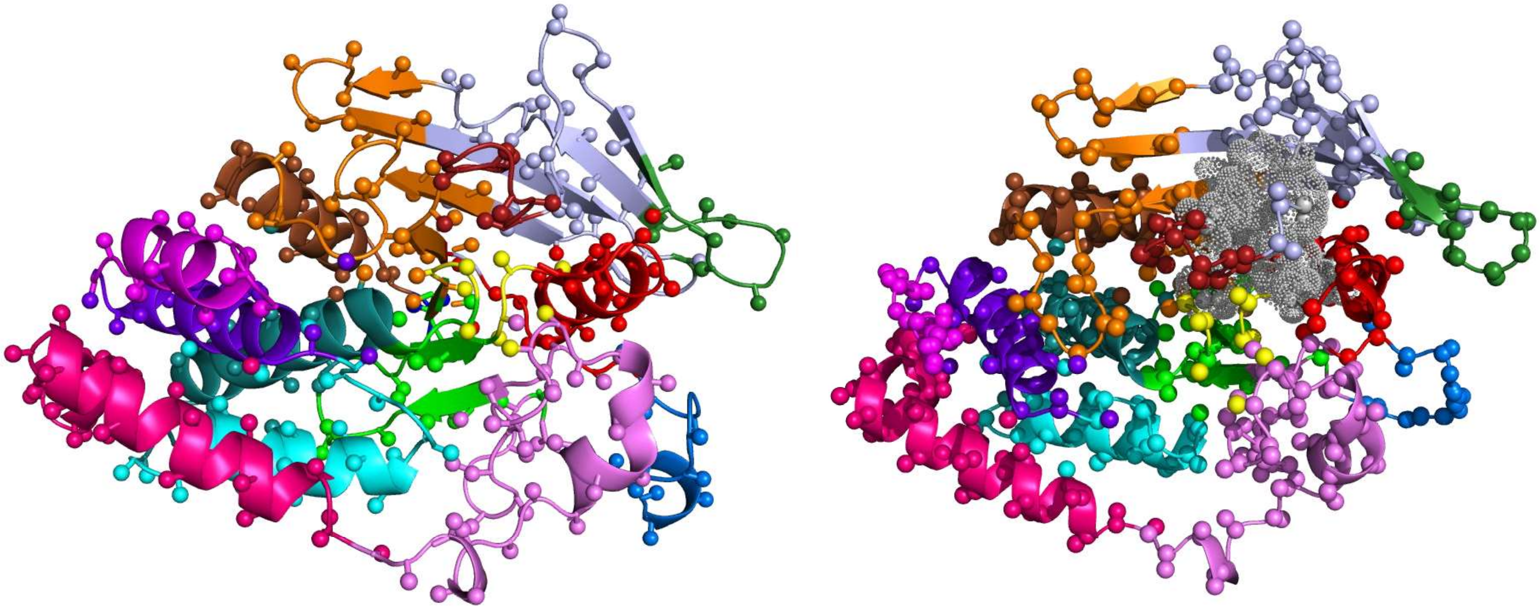

#### Supplementary Figure 20

**PTP1B apo**

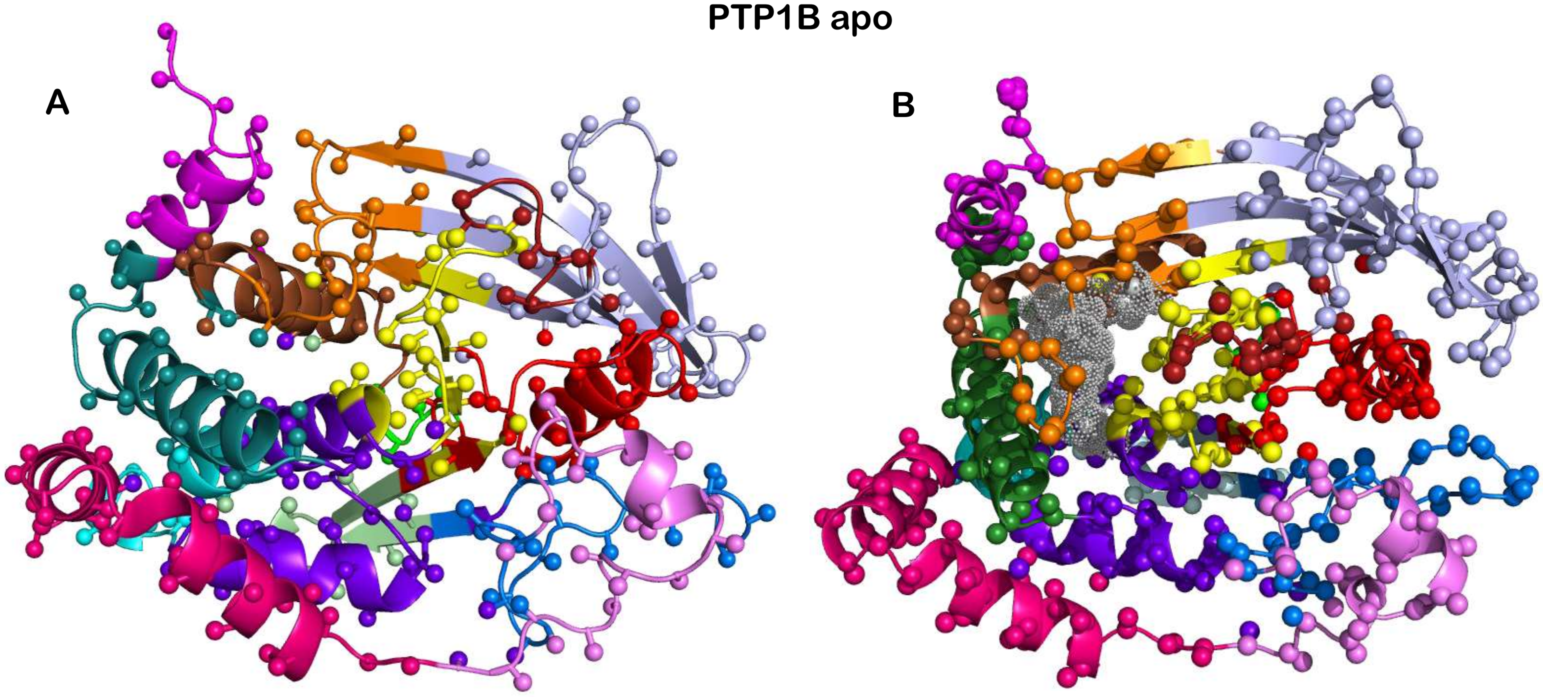


**PTP1B + pY**

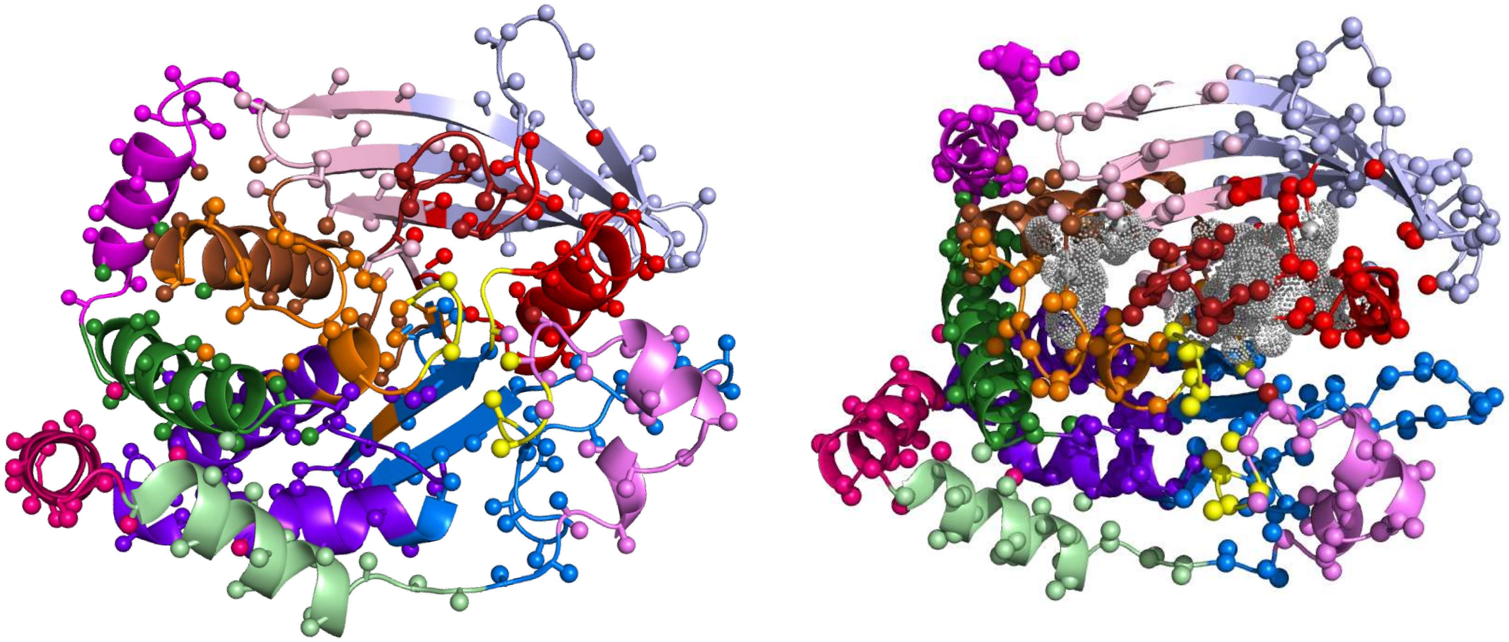

**Supplementary Figure 21**

**YopH apo**

**A**

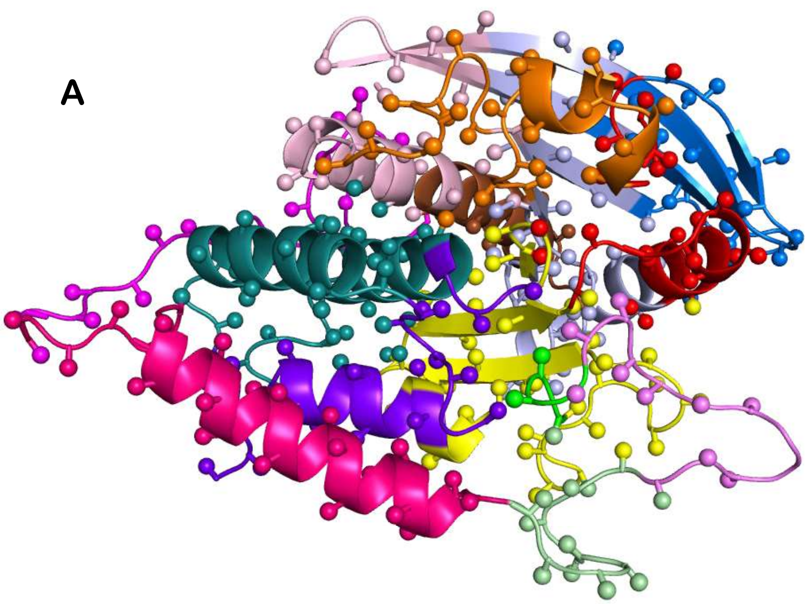

**B**

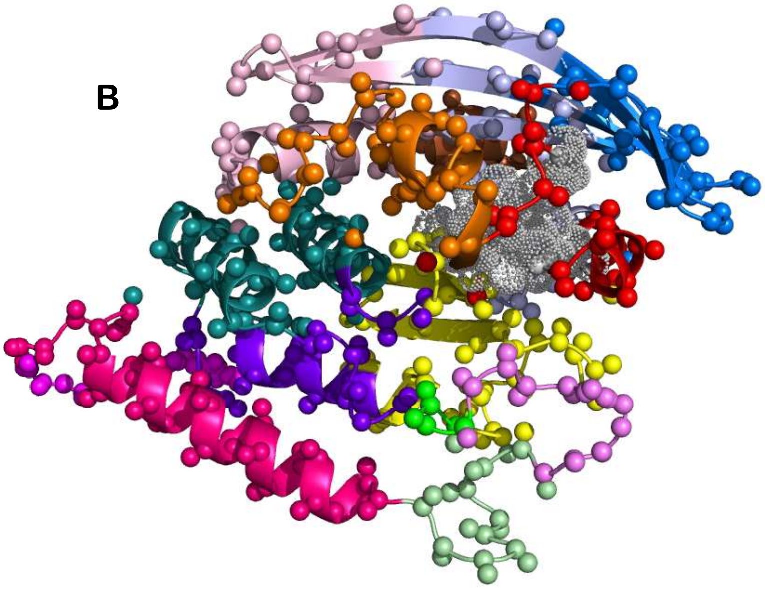

**YopH + pY**

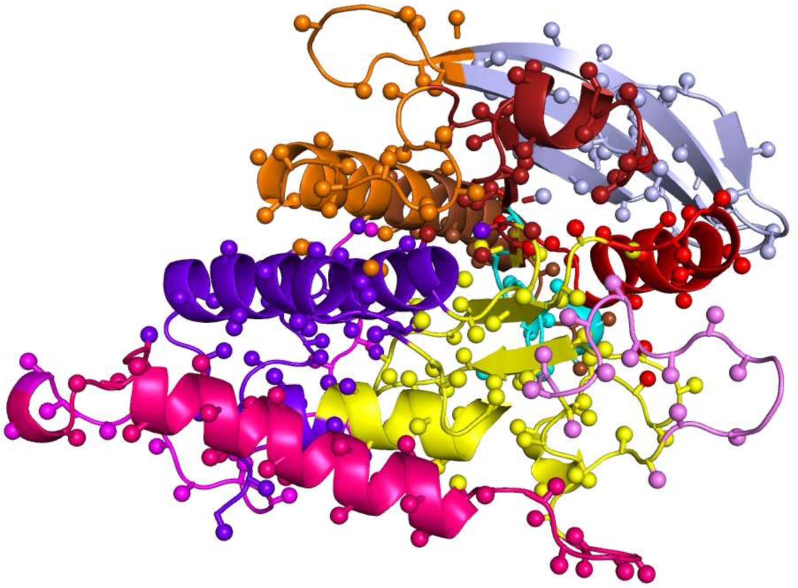

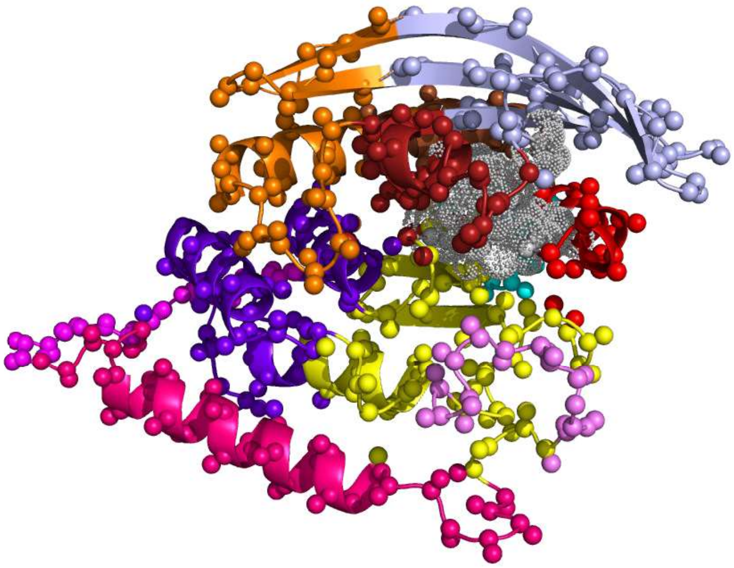

**Supplementary Figure 22**

**A**

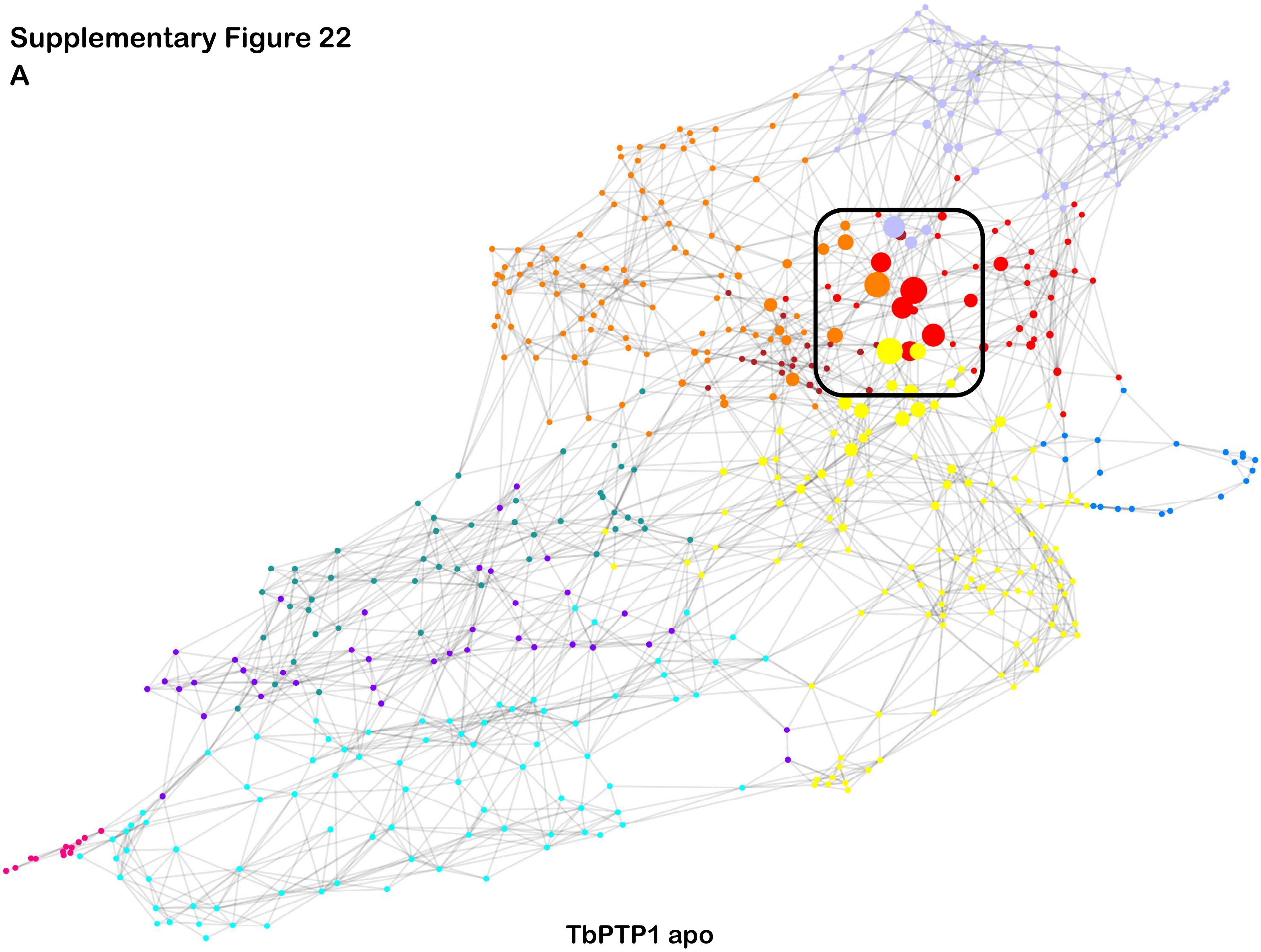

**TbPTP1 apo**

**B**

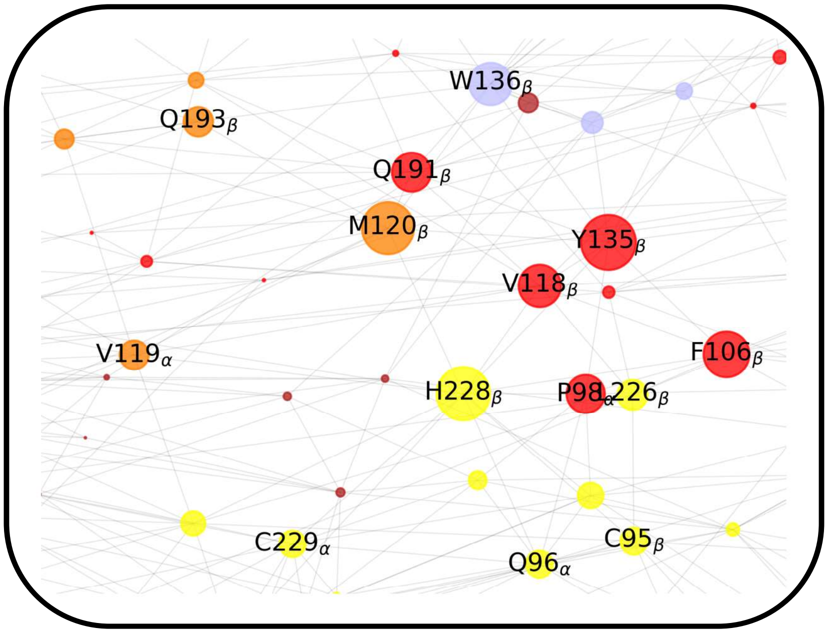

**Supplementary Figure 23**

**A**

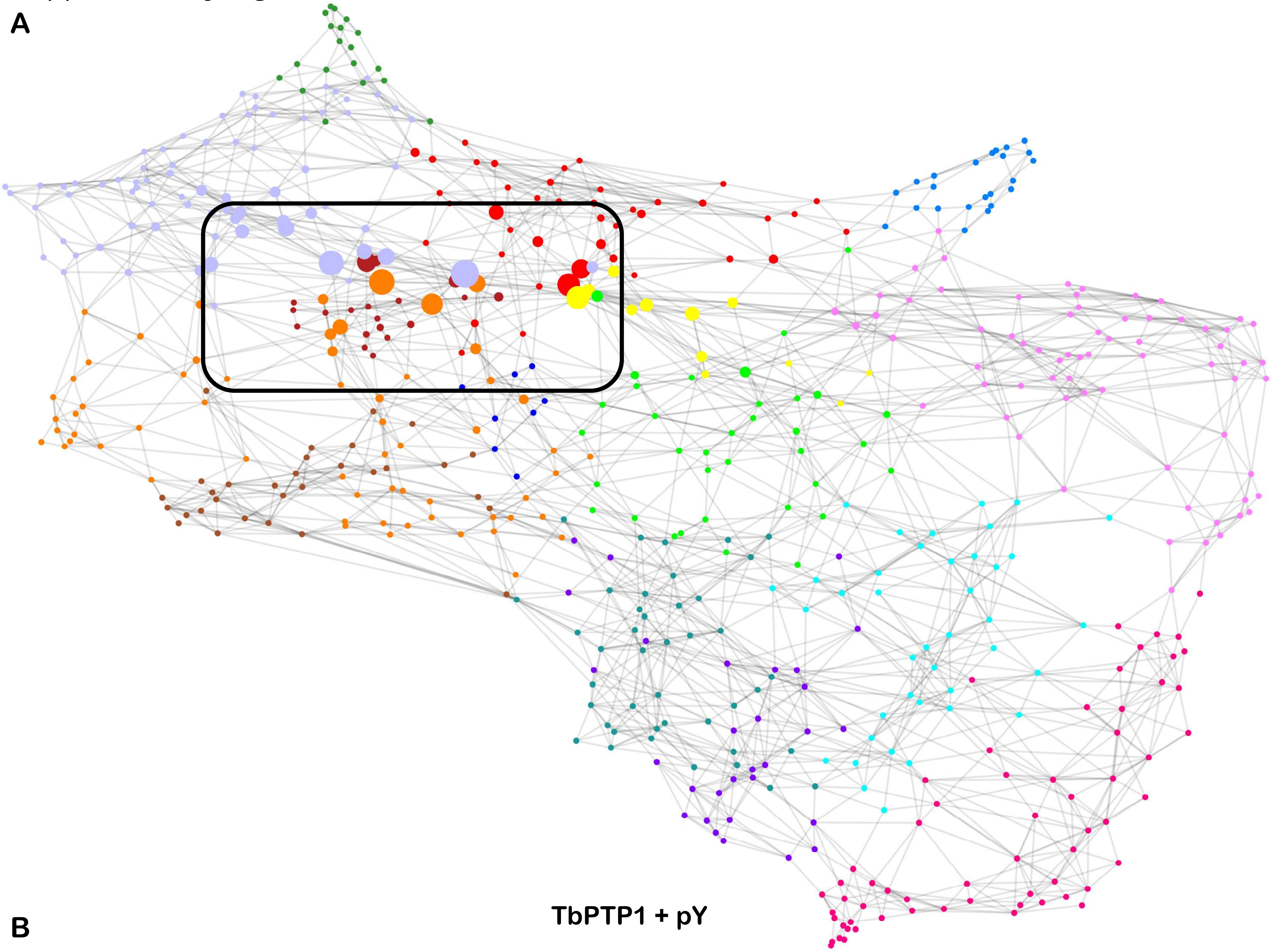

**B**

**TbPTP1 + pY**

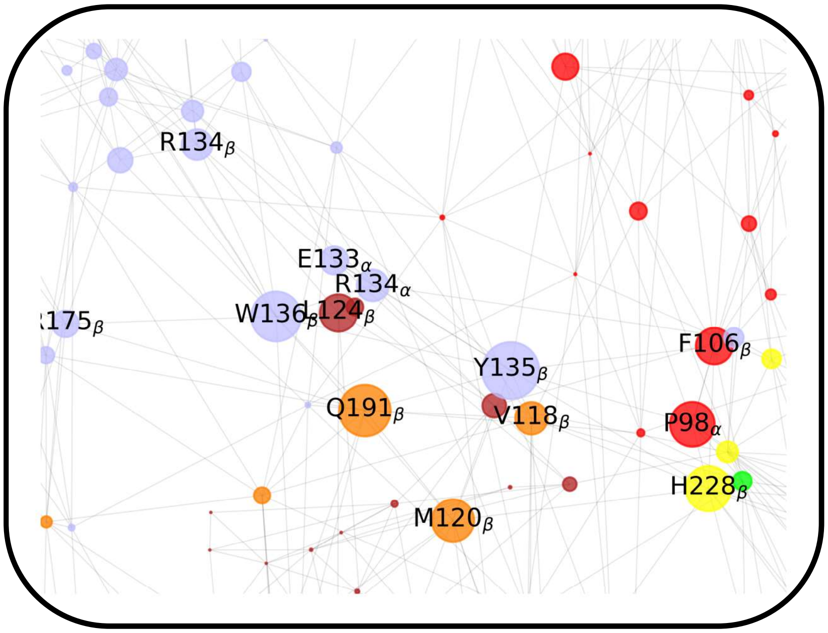

**Supplementary Figure 24**

**A**

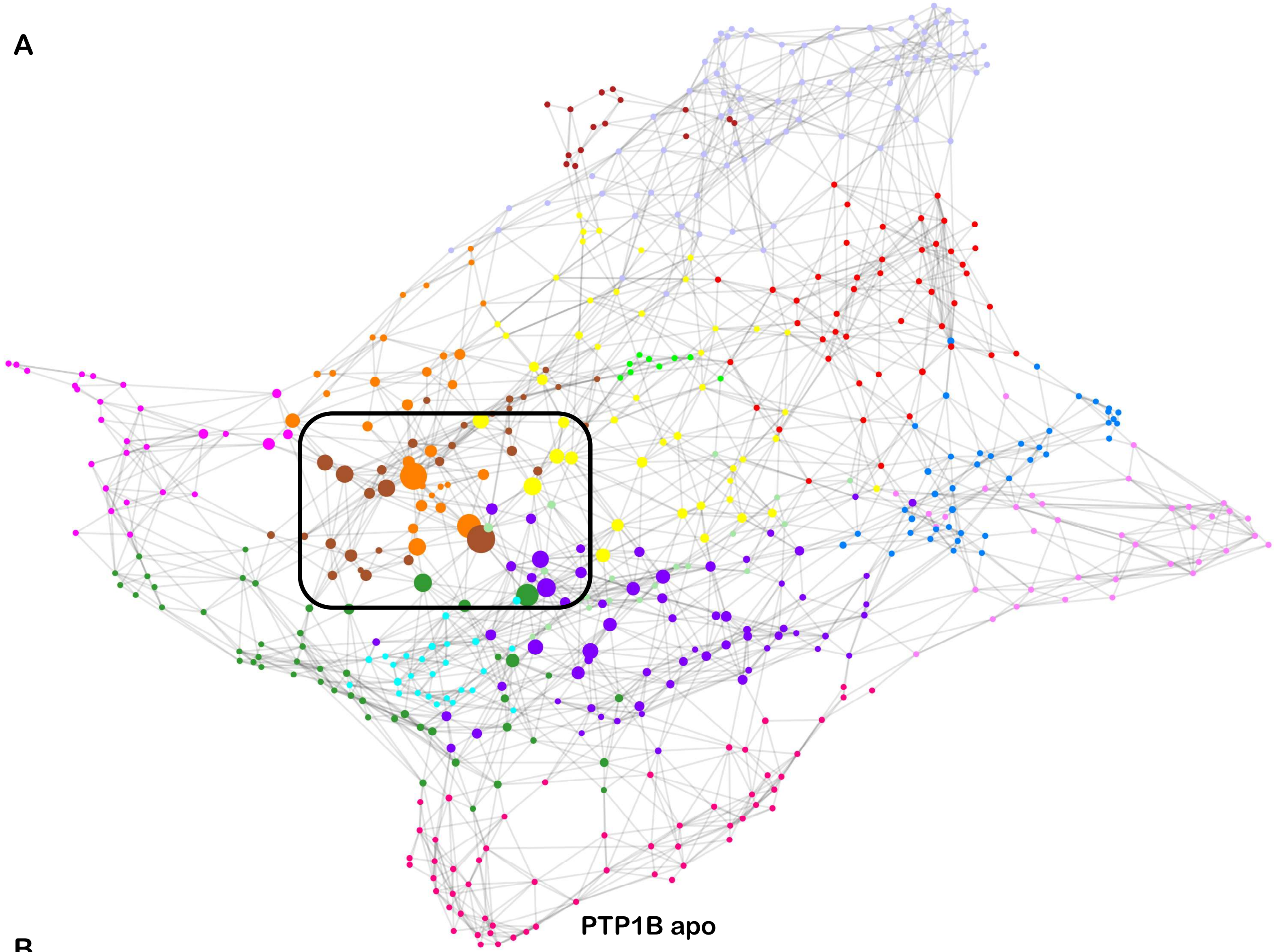

**B**

**PTP1B apo**

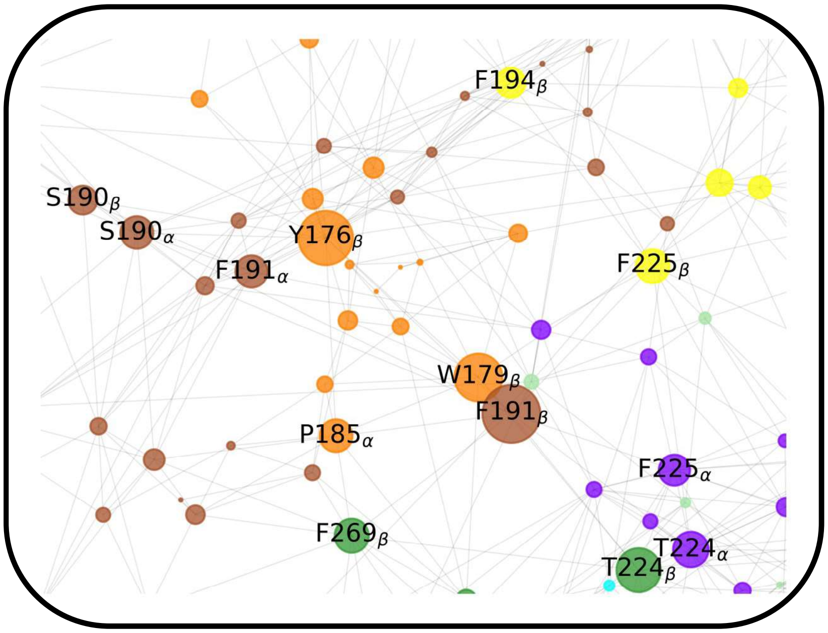

Supplementary Figure 25

A

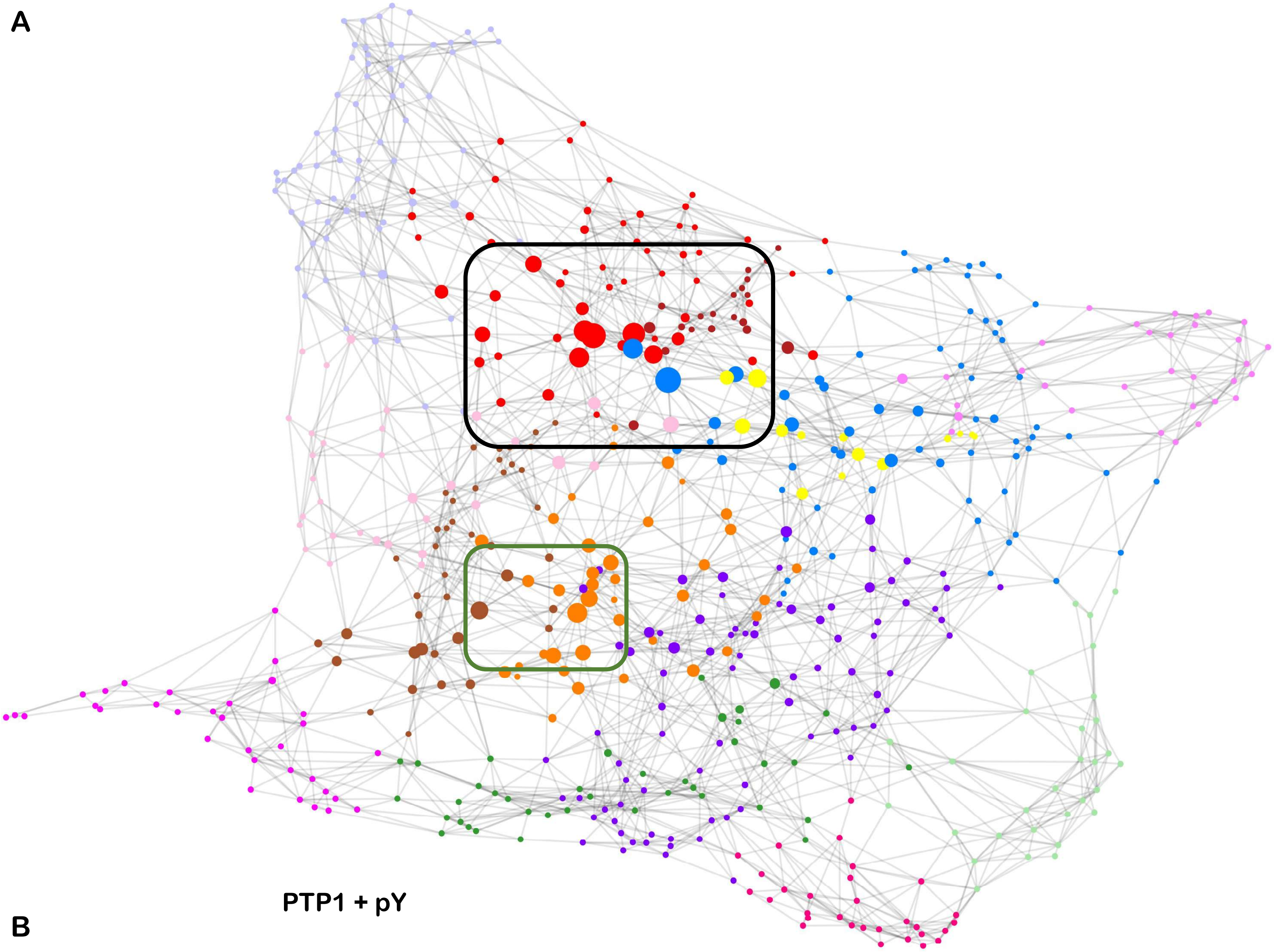


B

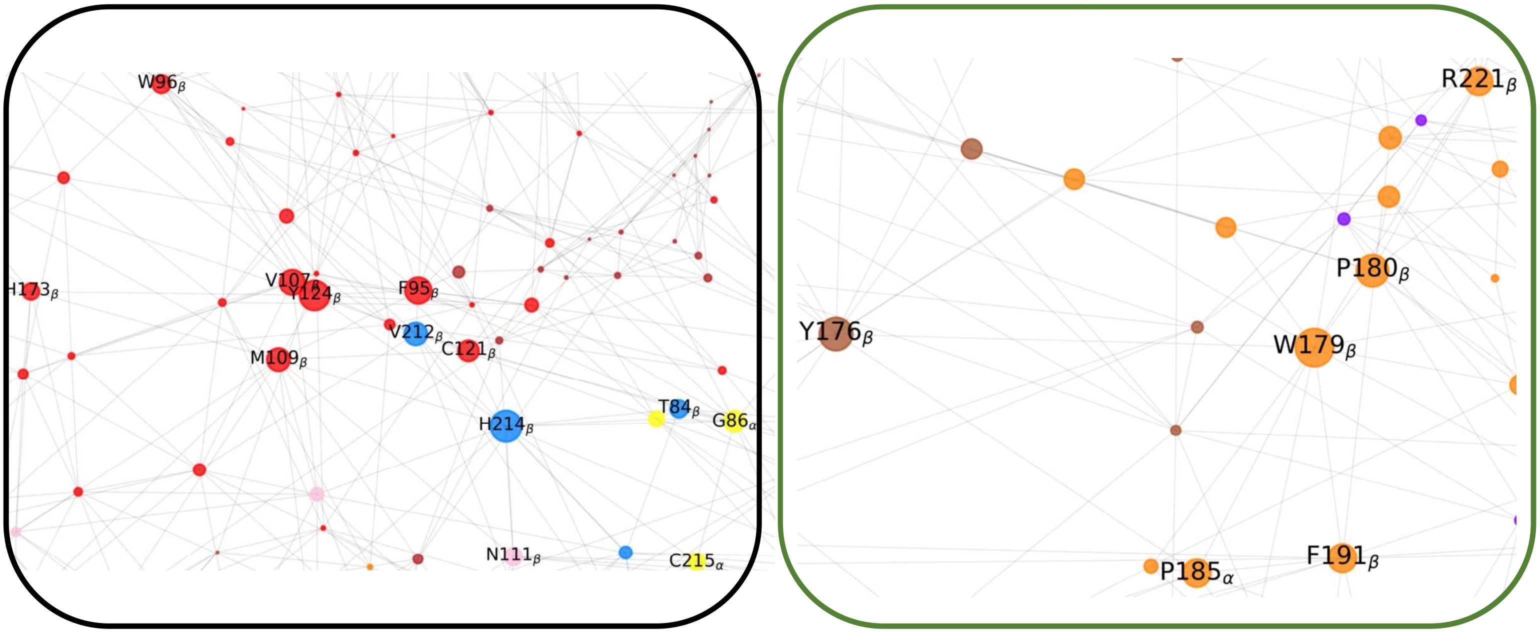

**Supplementary Figure 26**

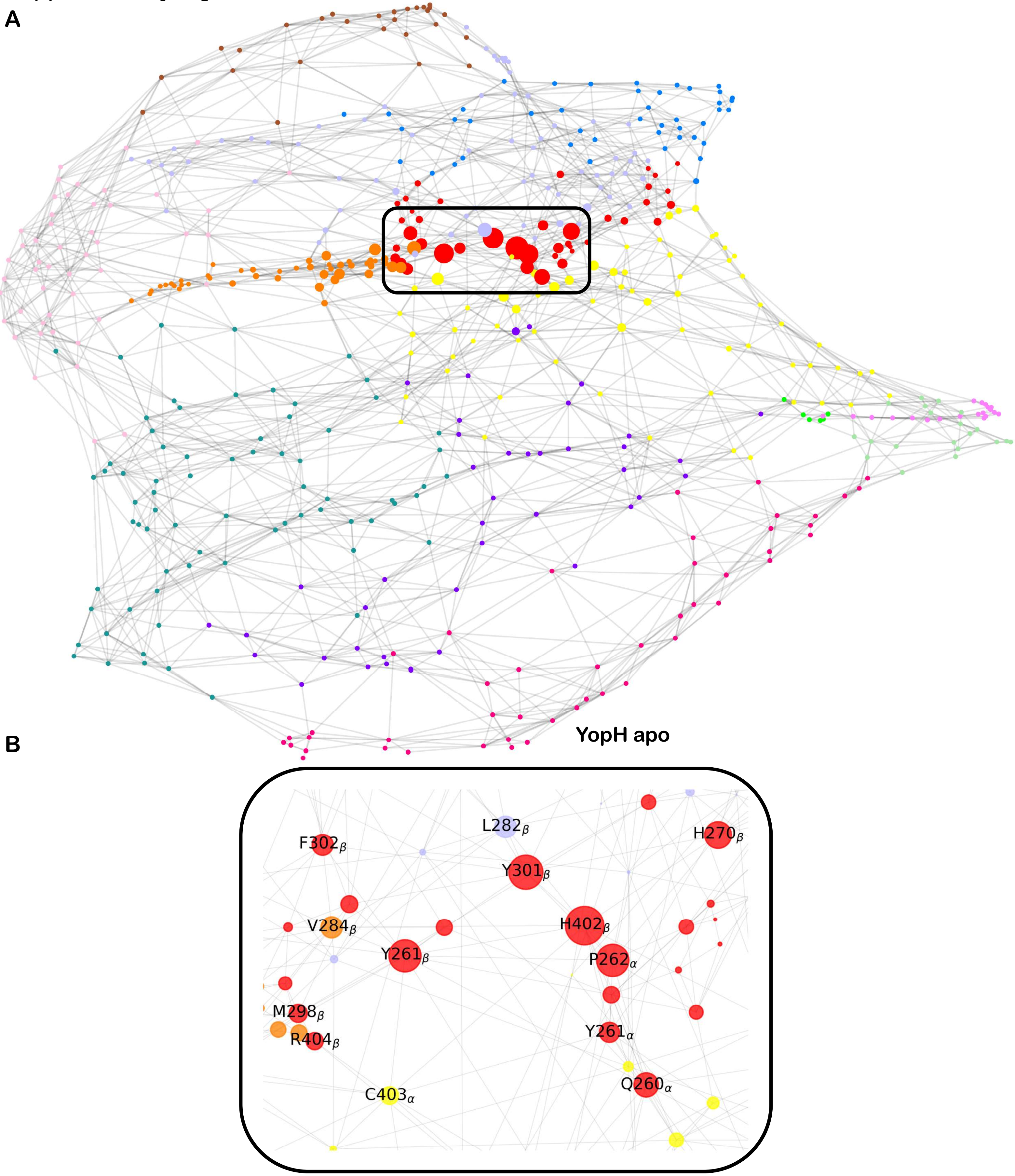

**Supplementary Figure 27**

**A**

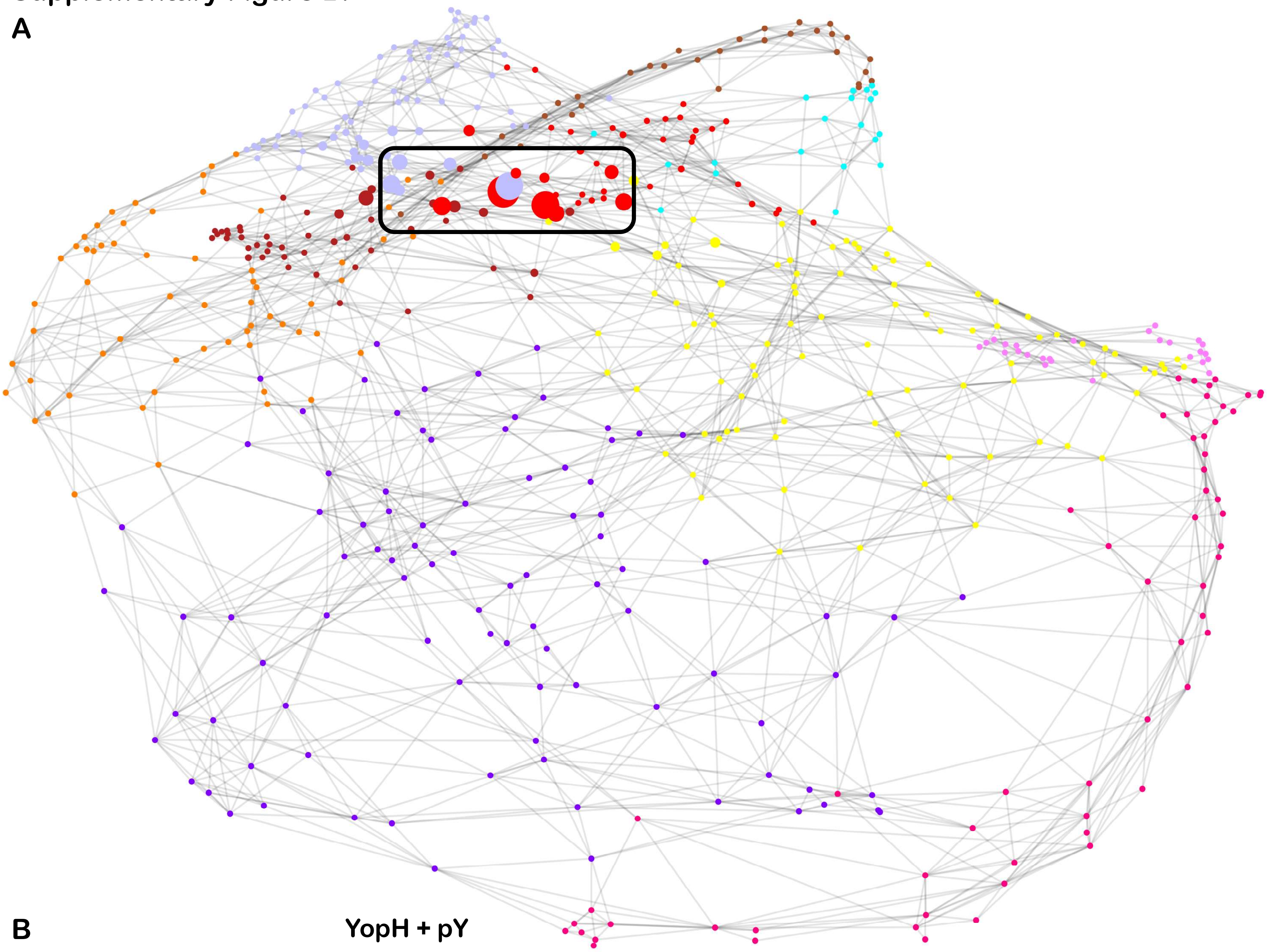

**B**

**YopH + pY**

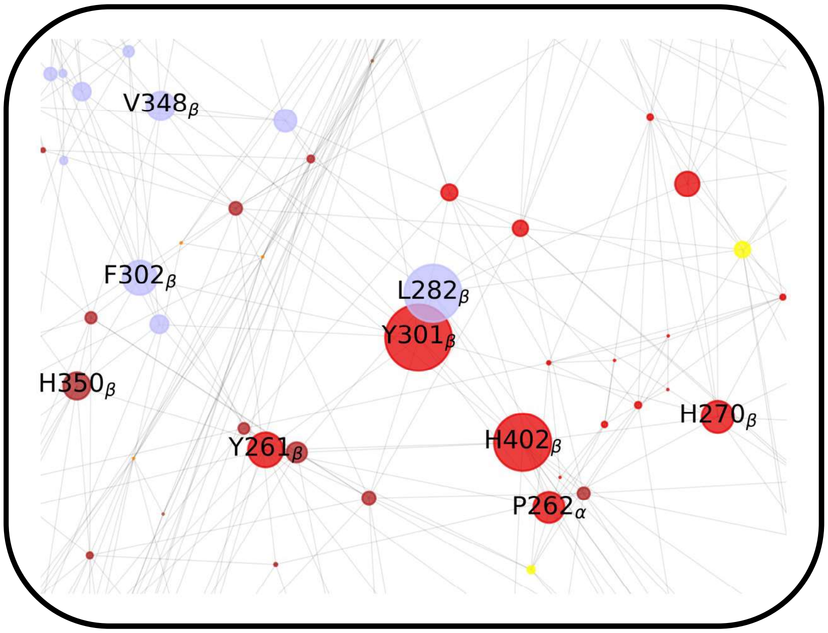
